# Deep axonal proteomics of human iPSC-derived neurons by microfluidic separation and DIA-MS

**DOI:** 10.64898/2026.07.17.739061

**Authors:** Clemens M Sauter, Zac Sandy, Milena Korneck, Vincent Albrecht, Melanie Kraft, Ramakrishnan Sivasubramanian, Barbora Kuttichova, Jared Sterneckert, Ludger Schöls, Alexandra Davies, Stefan Hauser

## Abstract

One of the defining features of neurons is their compartmentalisation into soma, dendrites and axon. Axons are highly specialised for the transmission of signals over long distances but also exhibit unique vulnerabilities in the context of neurodegeneration, implying a specialised axonal proteome. However, in-depth proteomic characterisation of axons has been limited by the difficulty of isolating pure axonal material in sufficient quantities for conventional mass spectrometry analysis.

Here, we combine microfluidic-based axon–soma separation with data-independent acquisition mass spectrometry (DIA-MS) on an Orbitrap Astral mass spectrometer for deep profiling of compartment-resolved proteomes of human induced pluripotent stem cell (iPSC)-derived cortical neurons. We quantify over 9,000 proteins in the somatodendritic compartment and ∼6,000 proteins in the axonal compartment, to our knowledge, representing the deepest human axonal proteome reported to date. Differential abundance analysis identifies 1,250 axon-enriched proteins with strong enrichment of axon-related pathways including vesicle-mediated transport, synaptic vesicle dynamics, and cytoskeletal organisation.

Extending this workflow to iPSC-derived lower motor neurons, we reveal a core set of 417 axon-enriched proteins shared between cortical and lower motor neurons, alongside subtype-specific axonal signatures whose functional differences become evident only through compartment-restricted analysis. We show broad coverage of genes for the neurodegenerative diseases hereditary spastic paraplegia and amyotrophic lateral sclerosis, establishing a quantitative reference for interpreting how disease-linked mutations may differentially affect axonal versus somatodendritic proteostasis.

This workflow and resource are readily applicable to other neuronal subtypes and disease models, paving the way for studying axonal proteome dynamics in health and disease.

**Highlights:**

- Deepest human axonal proteome with ∼6,000 proteins from iPSC-derived neurons
- Microfluidic separation paired with DIA mass spectrometry
- Axon-specific signatures masked in whole-cell analyses are revealed
- Core set of 417 axon-enriched proteins shared across neuron subtypes
- Framework for studying axon-selective vulnerability in neurodegeneration

**Graphical abstract:** 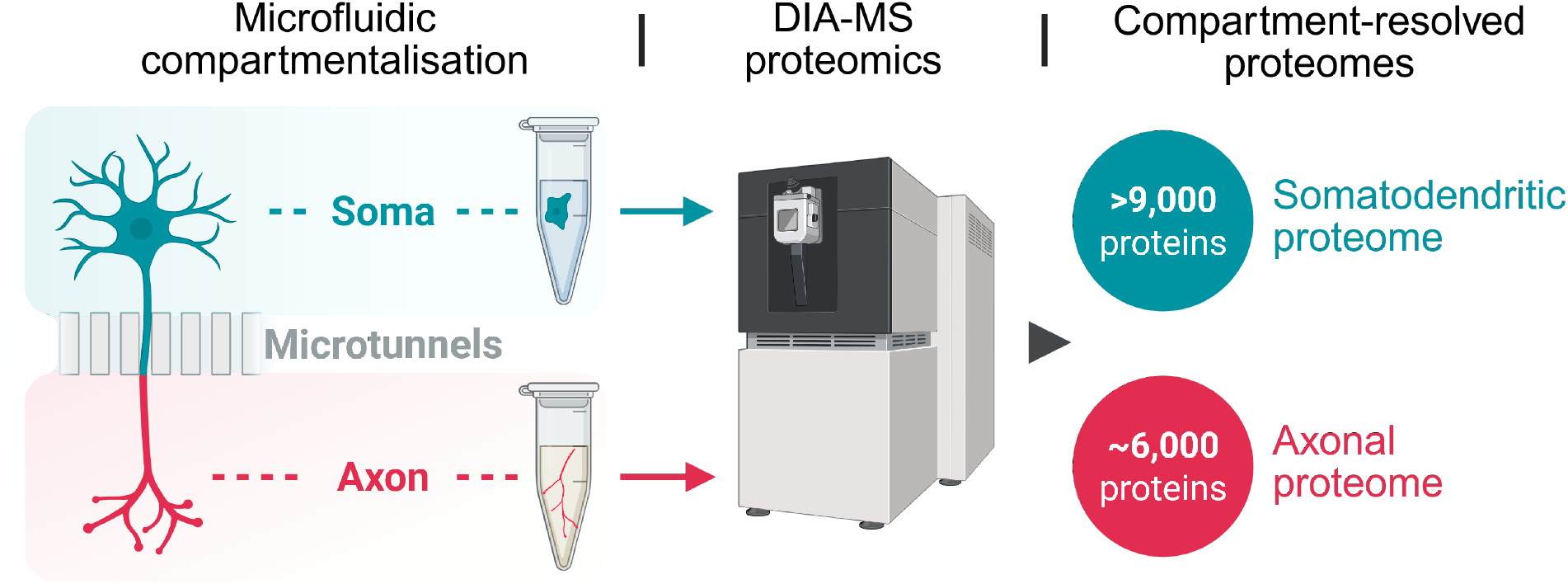

## Introduction

Neurons are highly polarised and structurally specialised cells, compartmentalised into soma (cell body), dendrites (for receiving signals) and axon (for transmitting signals). Motor neurons form the central circuitry of voluntary movement, with upper motor neurons in the cortex and lower motor neurons in the brainstem and spinal cord extending long axons to connect the central nervous system with muscles in the periphery. These axons can span more than a meter in humans, requiring efficient transport of organelles and proteins^1,2^, and local mechanisms of proteostasis^3,4^ to sustain function at sites distant from the soma. Their extraordinary length makes axons indispensable for neural connectivity but also renders them particularly susceptible to cellular stress and disruption.

This vulnerability is evident in motor neuron diseases, a group of neurodegenerative disorders that cause progressive damage predominantly to motor neurons. Hereditary spastic paraplegia (HSP) and amyotrophic lateral sclerosis (ALS) both feature progressive axonal dysfunction despite substantial genetic and clinical heterogeneity, with HSP predominantly affecting upper motor neurons^5,6^ whereas ALS affects both upper and lower motor neurons^7,8^. In HSP, evidence from patients and post-mortem data suggests that distal axons are often the first sites of pathology, preceding loss of neuronal cell bodies in a characteristic “dying-back” model of neurodegeneration^6,9^. Similarly, in ALS, motor neurons with their exceptionally long axons are preferentially affected^10,11^. However, the mechanisms driving these degenerative processes remain poorly understood, in part because the molecular landscape of axons is still incompletely defined.

Axons are increasingly recognised not as passive extensions of the soma but as compartments that sustain their own homeostasis through extensive trafficking, unique mitochondrial dynamics, and local protein synthesis^1–4,12,13^. Recent work suggests that axonal length is a key determinant of these spatially specialised processes, with distal regions exhibiting distinct molecular phenotypes in their cytoskeletal organisation, organelle distribution, mitochondrial content and calcium signalling^14^. Such compartment-specific mechanisms may represent critical points of vulnerability in motor neurons and underscore the importance of studying the specialised proteome of the axon.

Historically, direct proteomic analysis of axons has been challenging, largely because axons and cell bodies are densely intermingled in conventional neuronal cultures, making physical separation difficult. As a result, many recent attempts to define the axonal proteome have relied on proximity labelling followed by affinity purification and mass spectrometry^15,16^. While these techniques can robustly identify the most abundant axonal proteins, they often fail to detect lower abundance proteins and require stringent controls to account for relatively large labelling radii, which may capture proteins from neighbouring non-axonal compartments. Additionally, the use of a non-endogenous tag system can introduce artefacts such as overexpression, mislocalisation and altered protein-protein interaction networks, with potential downstream biological consequences.

Alternative approaches that aim to physically separate axons from cell bodies date to the introduction of Campenot chambers^17^ in the 1970s, with later improvements including membrane separation (e.g., Boyden chamber)^18–20^ and micropatterned substrate guidance systems^21^. However, these approaches suffer from inconsistent fluidic separation and dendrite contamination, limiting their use for pure axonal analysis. Consequently, microfluidic devices (MFDs) have emerged as the preferred method for axon-specific studies. MFDs consist of parallel chambers connected by micro-tunnels, which due to their length (typically 150–900 μm) permit only axons to grow through. This enables stringent compartment isolation, and integration with transcriptomic or image-based approaches has already proven successful in identifying locally translated axonal proteins and mechanisms underlying axon guidance^22,23^.

Despite these developments, MFDs have only recently been applied to proteomic profiling of human axons, achieving a limited depth of around 500 protein groups^24^. This is due to the low amounts of protein that can be harvested from the axonal compartment of MFDs (∼50 ng) and the challenges this presents for analysis with traditional data-dependent acquisition mass spectrometry (DDA-MS). Advances in data-independent acquisition mass spectrometry (DIA-MS) can substantially improve proteomic depth, with one study of microfluidically-isolated axons from primary mouse motor neurons identifying ∼2,400 protein groups using DIA-MS on an Orbitrap Exploris 480^25^. This represents a substantial improvement over the use of DDA-MS, but the introduction of the Orbitrap Astral mass spectrometer^26^ now promises further leaps in sensitivity and the potential to achieve unprecedented depth of analysis of the axonal proteome.

Here, we combined microfluidic compartmentalisation with high-sensitivity DIA-MS on an Orbitrap Astral to achieve deep, quantitative profiling of the axonal proteome of human induced pluripotent stem cell (iPSC)-derived cortical neurons, identifying ∼6,000 proteins in axonal compartments and ∼9,000 in somatodendritic compartments. This more than doubles the depth reported in previous axonal proteomics in either mice or human neurons^16,24,25^, establishing the most comprehensive map of the human axonal proteome to date. By extending this approach to iPSC-derived lower motor neurons, we define both conserved core and neuron subtype-specific axonal proteome signatures. These results demonstrate that deep, compartment-resolved proteomics can uncover both shared and specialised features of axonal biology, providing a quantitative framework for interrogating how disease-associated perturbations differentially impact axonal and somatodendritic compartments.

## Methods

### Experimental Design and Statistical Rationale

The study aimed to characterise compartment-resolved proteomes of human iPSC-derived neurons using microfluidic axon–soma separation coupled with data-independent acquisition mass spectrometry (DIA-MS). Samples comprised somatodendritic and axonal lysates from two independent cortical neuron iPSC lines (CN1, CN2) and one lower motor neuron iPSC line (MN1). Five biological replicates were independently cultured and harvested per condition, each replicate consisting of lysate pooled from two microfluidic devices to obtain sufficient material for proteomic analysis. Five replicates were selected to provide adequate statistical power for quantitative comparison while permitting exclusion of technical outliers that failed to cluster with other replicates by principal component analysis and hierarchical clustering; on this basis the fifth CN2 axonal replicate was excluded, giving final group sizes of CN1 (n = 5 axon, 5 soma), CN2 (n = 4 axon, 5 soma) and MN1 (n = 5 axon, 5 soma). Somatodendritic samples served as the biological control for axonal enrichment analysis, and the independent lines (CN1, CN2 and MN1) as biological controls for reproducibility and neuronal subtype-specific effects. All samples were processed using identical workflows. CN1 and CN2 samples were acquired first, with the two lines randomised with respect to each other within each compartment, with the CN somatodendritic block of samples being acquired before CN axonal samples. MN1 samples were subsequently acquired as a separate batch on the following day. No retention-time (RT) calibration peptides or other exogenous protein or peptide standards were spiked into samples prior to LC-MS analysis.

MS1 survey data were acquired for all samples. Precursors were sampled across 380–980 m/z using 150 non-overlapping DIA isolation windows of 4 Th width, with an average total cycle time of ∼1.6 s (full acquisition parameters are given in*LC-MS/MS analysis*). Ion mobility separation was performed using a FAIMS Pro device at a single compensation voltage (−40 V), no gas-phase fractionation across multiple compensation voltages or window sets was employed. For subsequent peptide detection, an empirical spectral library was generated from all biological replicates in this study (DIA-NN reanalysis mode) in combination with an in silico predicted library; library details are given in DIA data processing.

Differential abundance was assessed using two-sided Student’s t-tests with permutation-based false discovery rate (FDR) correction. The t-test was chosen because protein intensities were log₂ transformed and normalised prior to analysis, approximating a normal distribution, and permutation-based FDR was used to control for multiple testing across thousands of quantified proteins. Full statistical, imputation, clustering and enrichment parameters are described in*Quantification and statistical analysis*,*Fuzzy c-means clustering* and*Gene Ontology enrichment analysis*.

### Human iPSC lines and maintenance

Human iPSC lines used for differentiation into cortical neurons (CN1, CN2) were fully characterised in accordance with the scientific guidelines of the Human Pluripotent Stem Cell Registry (hPSCreg). The T12.9 WT FUS-eGFP reporter line (MN1), in which eGFP is fused to the C-terminus of endogenous FUS with a long linker, was generated as previously described^27^. NIL-iPSC lines were generated by PiggyBac transposition as previously described by De Santis et al^28^. All iPSC lines were maintained in Essential 8 (E8) medium (prepared in-house) on Matrigel-coated plates (Corning, cat# 354230) under standard conditions. Cells were passaged at 70–80% confluency and routinely tested for mycoplasma contamination by PCR.

### Cortical neuron differentiation (CN1 and CN2)

Cortical neurons were differentiated following a dual-SMAD inhibition protocol with minor modifications^29,30^. Briefly, iPSC colonies were expanded to ∼70-80% confluency, dissociated and replated onto Matrigel-coated plates in the presence of the ROCK inhibitor Y-27632 (10 µM; Selleckchem, cat# S1049) at a seeding density of approximately 3 × 10⁶ cells per well of a 6-well plate (Greiner Bio-One GmbH, cat# 657160). From days 0 to 7, medium was changed daily to 3N medium (1:1 DMEM/F-12 (Gibco, cat# 11320) and Neurobasal (Gibco, cat# 21103) supplemented with MEM NEAA (1x; Gibco, cat# 11140]) and B27 (1x; Gibco, cat# 17504-044)) containing SB-431542 (15 µM; Biozol, cat# SB431542) and LDN-193189 (500 nM; Sigma-Aldrich, cat# SML0559). On day 8, FGF2 (20 ng/mL; Preprotech, cat# 100-18B) was added. On day 9, neuroepithelial cells were passaged at a 1:3 ratio onto Matrigel-coated plates in 3N medium supplemented with FGF2 and Y-27632. FGF2 was gradually withdrawn between days 13-15, during which heparin (100 µg/mL, Sigma-Aldrich, cat# H3393-25KU) was added to the medium.

On day 19, cultures were cryopreserved in freezing medium (90% KnockOut Serum Replacement (Life Technologies, cat# 10828028), 10% DMSO (Sigma-Aldrich, cat# D4540), 10 µM Y-27632). Following thawing, cells were maintained in 3N medium with medium changes every other day. At day 26, cells were replated onto poly-L-ornithine (Sigma-Aldrich, cat# p8638)/Matrigel-coated substrates. To promote terminal neuronal differentiation, cultures were treated with 3N medium supplemented with small molecule inhibitors DAPT (10 µM; Sigma-Aldrich, cat# D5942) and PD0325901 (10 µM; Tocris Bioscience, cat# 4192) from days 27 to 29. Thereafter, cells were returned to standard 3N medium. By day 36, most neurons expressed the deep-layer cortical markers CTIP2 and TBR1. For long-term maturation, neurons were maintained in 3N medium supplemented with BDNF (10 ng/mL; Preprotech, cat# 450-02), GDNF (10 ng/mL; Preprotech, cat# 450-10-50UG), and NGF (10 ng/mL; Preprotech, cat# 450-01-20UG).

### Motor neuron differentiation (MN1)

Motor neurons were differentiated from doxycycline-inducible NIL (NGN2, ISL1, LHX3) iPSC lines^28^ using a staged medium protocol. On day 0, cells were dissociated and seeded onto Matrigel-coated plates at ∼5 × 10⁵ cells per well of a 6-well plate in E8 medium supplemented with Y-27632 (10 µM). On days 1 to 2, cultures were maintained in induction medium (DMEM/F-12 with MEM NEAA (1x) and doxycycline (1 µg/mL; Sigma-Aldrich, cat# D3072)). From days 3 to 5, medium was replaced daily with differentiation medium (Neurobasal with NEAA, B27 without vitamin A (1x; Gibco, cat# 12587010), doxycycline (1 µg/mL), DAPT (5 µM) and SU5402 (4 µM; Sigma-Aldrich, cat# SML0443).

On day 6, neuronal precursors were replated onto poly-L-ornithine/Matrigel-coated plates in maturation medium (Neurobasal with B27 without vitamin A, BDNF (20 ng/mL), GDNF (10 ng/mL), ascorbic acid (0.2 mM; Tocris, cat# 4055) and Y-27632 at 10 µM). Neurons were maintained in maturation medium with changes every 2-3 days. After ≥7 days of maturation, neurons expressed canonical motor neuron markers including HB9, ISL1, ChAT and SMI-32.

### Microfluidic culture

Microfluidic XonaChips (Xona Microfluidics, cat# XC900) were preconditioned with XC Pre-Coat and sequentially coated with poly-L-ornithine followed by Matrigel (1:45 dilution, 1 h, 37°C).

For cortical cultures (CN1 and CN2), neurons were dissociated and loaded into microfluidic devices at day 26 of differentiation. For motor neuron cultures (MN1), cells were dissociated and loaded at day 6 of differentiation. In all cases, cells were filtered through a 30 µm cell strainer and resuspended in the respective culture medium supplemented with ROCK inhibitor Y-27632 (10 µM).

Approximately 3 × 10⁵ neurons were loaded into the somatic compartment in ∼5 µL, allowed to settle for 30 min, and then overlaid with the corresponding maintenance medium containing ROCK inhibitor Y-27632 (10 µM). Cultures were maintained according to their respective differentiation schedules. Cultures were maintained until day 58 of differentiation prior to lysis, sample collection, or fixation for immunocytochemistry.

### Immunocytochemistry

For validation of neuronal identity and compartmentalization, immunocytochemistry was performed on neurons cultured in microfluidic XonaChips as well as on parallel cultures maintained in standard 96-well plates (PhenoPlate, cat# 6055302). Cultures were washed once with PBS (Gibco, cat# 14190144) and fixed with 4% paraformaldehyde (Thermo Fisher Scientific, cat# J19943.K2) for 15 minutes at 37°C, followed by three washes with PBS. Fixed samples were permeabilized and blocked simultaneously in PBS containing 0.1% Triton X-100 (Sigma-Aldrich, cat# 9036-19-5) and 5% bovine serum albumin (BSA; Roth, cat# T844.4) for 1 hour at room temperature.

Primary antibodies were diluted in PBS-T with 5% BSA and incubated overnight at 4°C. The following primary antibodies were used: anti-MAP2 (1:5,000, chicken polyclonal; BioLegend, cat# 822501), anti-βIII-tubulin (1:1,000, mouse monoclonal; Sigma-Aldrich, cat# T8660), anti-CTIP2 (1:500, rat monoclonal; Abcam, cat# ab18465), anti-TBR1 (1:500, rabbit monoclonal [recombinant], Abcam, cat# ab183032), anti-ChAT (1:200, goat; Milipore, cat# AB144P), anti-HB9 (1:100, rabbit; Abcam, cat# ab221884). Synaptophysin-1 was detected using an Alexa Fluor 488-conjugated anti-Synaptophysin antibody (1:500, mouse monoclonal; Proteintech, cat# CL488-67864).

After primary antibody incubation, samples were washed three times with PBS-T (5 minutes per wash). Species-appropriate Alexa Fluor-conjugated secondary antibodies were diluted 1:1,000 in PBS-T and applied for 1 hour at room temperature, protected from light. Samples were washed three times with PBS- T. Nuclei were counterstained with DAPI (1 µg/mL) diluted in PBS-T for 30 minutes at room temperature, followed by three final washes with PBS.

Fluorescent images were acquired using a Nikon laser-scanning confocal microscope with a motorized stage. Large-area images of entire microfluidic devices were obtained by automated tile scanning and stitching. Z-stacks were collected to account for focal and height differences across device regions. Imaging settings were kept constant across conditions.

Stitching and initial processing were performed in Nikon NIS-Elements AR (v 6.10.02). Maximum intensity projections were generated where appropriate and Fiji (ImageJ)^31^ was used for consistent brightness and contrast adjustments across fluorescence channels and experimental conditions.

### Sample collection and lysis

At day 58, axonal and somatic compartments were harvested from microfluidic devices. Per biological replicate, lysates from two microfluidic devices were pooled. Low-protein-binding tubes were used throughout. The axonal compartment was washed twice with 200 µL pre-warmed PBS. After aspiration, 10 µL iST-LYSE buffer (PreOmics GmbH, cat# NC1699276) was added to the upper axonal well while maintaining a larger volume of PBS on the somatodendritic side to prevent backflow across the microtunnels. The solution was pipetted up and down five times and allowed to flow to the lower axonal well. Pooled lysates were immediately heated at 96°C for 5 minutes with shaking at 1,400 rpm to inactivate proteases. Samples were briefly cooled on ice, snap-frozen in liquid nitrogen and stored at -80°C. The somatodendritic compartment was processed in the same way, except that 40 µL lysis buffer was used per device.

### Proteolytic digestion and peptide cleanup

Frozen lysates were thawed and heated at 96°C for 5 minutes, followed by sonication in a Bioruptor (15 cycles, 30 seconds on/off, maximum intensity). Protein concentrations of somatodendritic samples were determined using a tryptophan fluorescence assay. For somatodendritic samples, 50 µg protein were digested with 1 µg trypsin and 1 µg LysC overnight at 37°C with shaking at 1,200 rpm. Protein concentrations of axon samples were too low to be directly quantified and were estimated to be ∼50 ng based on preliminary mass spectrometry experiments. For axonal samples, the entire lysate was digested with 0.02 µg trypsin and 0.02 µg LysC under the same conditions.

Peptides were acidified with isopropanol/1% TFA (5× sample volume) and desalted using poly(styrenedivinylbenzene) reverse-phase sulfonate (SDB-RPS) stage-tips, as previously described^32^. Stage-tips were activated with 100 µL acetonitrile, equilibrated with 100 µL 30% methanol/1% TFA followed by 100 µL 0.2% TFA. After sample loading (20 µg for soma, entire volume for axon), stage-tips were washed with 100 µL isopropanol/1% TFA and 100 µL 0.2% TFA, then eluted with 60 µL 80% acetonitrile/1.25% ammonium hydroxide. Eluates were dried by vacuum centrifugation and stored at−20°C. Somatodendritic samples were resuspended in buffer A (0.1% formic acid) to a final concentration of 125 ng/µL.

For LC-MS analysis, peptides were loaded onto Evotip C18 trap columns (Evosep Biosystems; cat# EV2001), according to the manufacturer’s instructions. Evotips were activated in 1-propanol, washed with 50 µL buffer B (0.1% formic acid in acetonitrile), soaked again in 1-propanol and equilibrated with buffer A. Samples were loaded (300 ng for soma, entire sample for axon), washed with buffer A and maintained with 150 µL buffer A to prevent drying.

### LC-MS/MS analysis

Samples were analysed using an Evosep One liquid chromatography system (Evosep Biosystems) coupled to an Orbitrap Astral mass spectrometer (Thermo Fisher Scientific) operated in data-independent acquisition (DIA) mode. Peptides were separated over 21 minutes on an 8 cm, 150 µm ID Aurora column (IonOpticks) using the 60 samples per day (SPD) method. The Orbitrap Astral was interfaced with a FAIMS Pro device operated with a compensation voltage of −40 V. The spray voltage was set to 1900 V.

MS1 scans (380–980 m/z) were acquired in the Orbitrap at a resolution of 120,000 with an AGC target of 500% and a maximum injection time of 3 ms. DIA scans were acquired in the Astral analyzer covering a precursor mass range of 380–980 m/z using 150 windows of 4 Th width. Fragment ions were detected over a scan range of 150–2000 m/z with a maximum injection time of 8 ms and an AGC target of 500%. HCD fragmentation was performed with 25% normalized collision energy.

### DIA data processing

Raw data were analysed using DIA-NN (version 1.8.1)^33^. An in silico spectral library was predicted directly from FASTA sequences in library-free mode, using the human Uniprot Swiss-Prot reference proteome (canonical and isoform sequences, downloaded 16 January 2025, available: https://www.uniprot.org/proteomes/UP000005640) combined with a common proteomics contaminants database^34^. Library prediction was performed with peptide lengths of 7–30 residues, precursor charge states 2–4, precursor m/z 300–1800, fragment m/z 200–1800, a 1% precursor-level FDR. The input library comprised 4,947,825 precursors corresponding to 42,860 protein isoforms (20,629 proteins, 20,252 genes after reannotation to the sequence database). Identical library and search settings were applied across four separate analyses (all samples combined, CN-only, axon-only, and soma-only), the parameters below apply to all four.

N-terminal methionine excision was enabled with trypsin specificity (cleavage after K/R) and a maximum of one missed cleavage. Oxidation of methionine was set as variable modification and cysteine carbamidomethylation as a fixed modification. Mass accuracy was fixed at 5 ppm for MS1 and 10 ppm for MS2, with a scan window radius of 6. The reanalysis mode (--reanalyse) was enabled, generating an empirical spectral library from the DIA data in the first pass which was used to requantify all runs in the second pass, with this empirical library filtered at 1% precursor and protein-group FDR. This procedure was applied independently within each of the four analyses. Relaxed protein inference (--relaxed-prot-inf) was applied and quantification used RT profiling with fixed-width peak centre integration; interference removal from fragment elution curves was disabled. The predicted spectral library included predicted retention times and ion mobility was not used as a library dimension as FAIMS was operated at a single compensation voltage (−40 V). Decoys were generated internally by DIA-NN’s target–decoy approach, and protein-group q-values were computed using proteotypic peptides only. Library-level FDR does not apply to the in silico predicted input library; the empirical libraries generated during reanalysis were filtered at 1% FDR as noted. The final output of each analysis was filtered at 1% FDR at both precursor and protein-group levels.

### Quantification and statistical analysis

Protein quantification was derived from DIA-NN output. Isoforms within a protein group were collapsed into a single representative, retaining only the isoform with the greatest summed peptide count across all samples for downstream analysis. For axon-only and soma-only datasets, MaxLFQ-normalised protein intensities generated by DIA-NN were used directly. For combined axon–soma analyses, raw (un-normalised) DIA-NN protein-group quantities (the PG.Quantity column of the report.tsv file, aggregated from precursor-level quantities rather than via the MaxLFQ algorithm) were median-normalised across samples to account for global differences in intensity distributions between axonal and somatodendritic proteomes that may violate assumptions underlying MaxLFQ-based normalisation. Single-peptide protein identifications are included in quantitative analyses and gene ontology enrichments but are not highlighted as individual protein-level findings. Proteins identified by multiple distinct peptides comprise the majority of reported identifications (>87% in all datasets).

All intensities were log₂ transformed prior to statistical analysis. Proteins were annotated and filtered based on data completeness. For axon–soma datasets, proteins were required to have valid values in at least 70% of samples within at least one experimental group (soma-CN1, soma-MN1, axon-CN1, or axon-MN1). For axon-only or soma-only analyses, proteins were required to be quantified in at least 70% of samples within at least one condition (CN1 or MN1).

Missing values were imputed in Perseus (v2.1.3)^35^ using a normal distribution-based approach (width = 0.3, downshift = 1.8), applied separately to each sample column.

Differential protein abundance was assessed using two-sided Student’s t-tests with permutation-based false discovery rate (FDR) correction (FDR < 0.05; 999 permutations). For both axon–soma and CN1 versus MN1 comparisons, an S₀ parameter^36^ of 0.1 was applied. For the latter, an additional log₂ fold-change threshold > 1 was imposed.

Principal component analysis was performed on centered, non-scaled log₂-transformed protein intensities using prcomp in R (version 4.4.2). Plots were generated in R using ggplot2.

### Fuzzy c-means clustering

Protein abundances from shared axonal proteins across CN1 and MN1 replicates were z-score normalized prior to clustering. Fuzzy c-means clustering^37^ was performed independently on the axonal proteome (5,851 proteins) and the somatodendritic proteome (9,717 proteins) using the Mfuzz package^38^ (version 2.66.0) in R (version 4.4.2). The fuzzifier parameter was estimated using the mestimate function and proteins were partitioned into three clusters in each dataset. Core cluster members were defined as those with a maximum membership score ≥ 0.7. Hierarchical clustering of samples was performed using complete linkage with Euclidean distance. Clusters were assigned biological labels (Shared, CN1-enriched, MN1-enriched) based on the difference between mean centroid expression values for CN1 and MN1 samples. One MN1 axonal replicate (MN1 Axon 5) and one MN1 somatodendritic replicate (MN1 Soma 4) were identified as outliers based on their clear separation from respective group clusters in PCA space and were excluded from the clustering analysis. These exclusions were applied only to the fuzzy c-means clustering analyses; all other comparisons retained the full replicate set.

### Gene Ontology enrichment analysis

For gene ontology (GO) analysis of axon and somatodendritic enriched proteins (Fig. 3, S5), WebGestalt 2024^39^ was used with weighted set cover redundancy removal and a Benjamini-Hochberg-corrected FDR threshold of < 0.05 FDR. A custom background comprising all proteins identified across all replicates and conditions was applied. Gene Ontology enrichment analysis of mFuzz clustered protein groups was performed in R using the clusterProfiler package (version 4.14.4;^40^) with annotation from org.Hs.eg.db. Core cluster members (membership ≥ 0.7) from the CN1-enriched and MN1-enriched clusters were tested for enrichment of GO Cellular Component (GOCC), GO Biological Process (GOBP) and KEGG pathway terms against a background set comprising all proteins detected in the respective dataset. Terms with a Benjamini-Hochberg-adjusted p-value < 0.05 were considered significantly enriched.

### Data availability

The mass spectrometry proteomics data, spectral library and FASTA files and DIA-NN analysis output (including precursor and protein-level report matrices) used for analyses have been deposited to the ProteomeXchange consortium via PRIDE and are available with identifier PXD080544.

## Results

### Microfluidic compartmentalisation of iPSC-derived cortical neurons

To enable the selective analysis of axonal proteomes, we established a microfluidic workflow using commercially available microfluidic devices (MFDs) that physically separate axons from somata and dendrites. The MFDs consist of a proximal somatodendritic compartment, where neurons are seeded, connected to a distal axonal compartment via microtunnels (900 µm long, 7 µm wide) that permit axonal but not somatodendritic passage (Fig. 1A). Human iPSC-derived cortical neurons were seeded in MFDs and axons extended through the microtunnels over four weeks, densely populating the distal compartment and enabling independent harvest of somatodendritic and axonal compartments for proteomic profiling. Although we refer to somatodendritic and axonal compartments throughout the text, the somatodendritic compartment contains axons in addition to somata and dendrites and should therefore be considered somatodendritic-enriched relative to the axon-pure compartment. Immunocytochemistry confirmed effective compartmentalisation, with nuclear (DAPI) and dendritic (MAP2) markers largely confined to the somatodendritic compartment whereas the cytoskeletal marker (βIII-tubulin) was present throughout, consistent with axonal projections extending through the microtunnels (Fig. 1B). Synaptophysin-1 was also detectable in both compartments, reflecting the presence of presynaptic vesicles in axon terminals and confirming axon maturity (Supplemental Fig. S1).

**Fig. 1.**
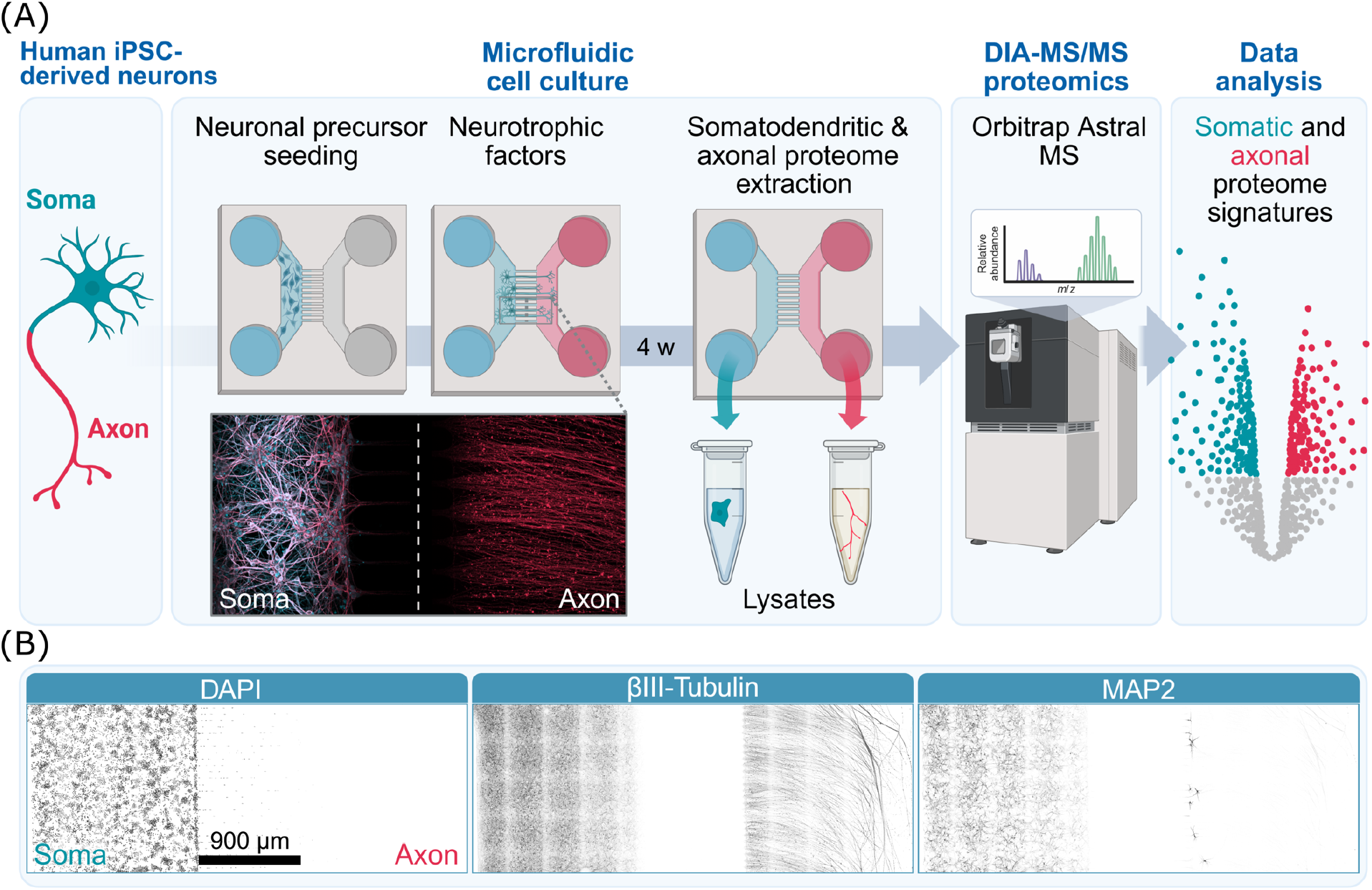
Microfluidic data independent acquisition mass spectrometry (DIA-MS) workflow. **A**, experimental workflow integrating microfluidic axon–soma separation with DIA-based proteomics. Human iPSC-derived neurons are seeded into the proximal somatodendritic (soma) compartment of XONA microfluidic devices with 900 µm length microtunnels. Axons extend through the microtunnels into the distal axonal (axon) chamber under neurotrophic cues, allowing selective collection of somatodendritic-enriched and axonal lysates, followed by proteomic profiling using DIA-MS on an Orbitrap Astral mass spectrometer. **B**, immunocytochemistry (ICC) validating compartment separation. DAPI and MAP2 signals were restricted to the somatodendritic (Soma) chamber; βIII-tubulin was detected in both (Soma and Axon) compartments. Synaptophysin-1 (SYP1) was present in both compartments (see Supplemental Fig. S1).

### Orbitrap Astral DIA-MS enables deep analysis of the axonal proteome

While MFDs were effective at isolating somatodendritic and axonal compartments, total peptide yield from two pooled axonal compartments after digest was estimated to be only ∼50 ng. To maximise the achievable depth of analysis from this limited amount of starting material, we optimised a high sensitivity DIA-MS method on an Orbitrap Astral mass spectrometer using 50 ng of peptide input (Supplemental Fig. S2). We then applied the optimised method to analyse the proteomes of somatodendritic and axonal compartments harvested from MFD cultures of two independent iPSC-derived cortical neuron lines (CN1 and CN2; 4 or 5 replicates per line, each pooled from 2 MFDs). Using a 21-minute liquid chromatography (LC) gradient, we quantified a total of 9,676 proteins in somatodendritic samples and 7,871 in axonal samples at 1% false discovery rate (FDR), with an average of 9,382 and 6,029 proteins per replicate respectively (CN1 vs CN2: 9,376 vs 9,389 for soma; 6,564 vs 5,359 for axon), after filtering out common contaminants and low-confidence isoform identifications (Fig. 2A).

**Fig. 2.**
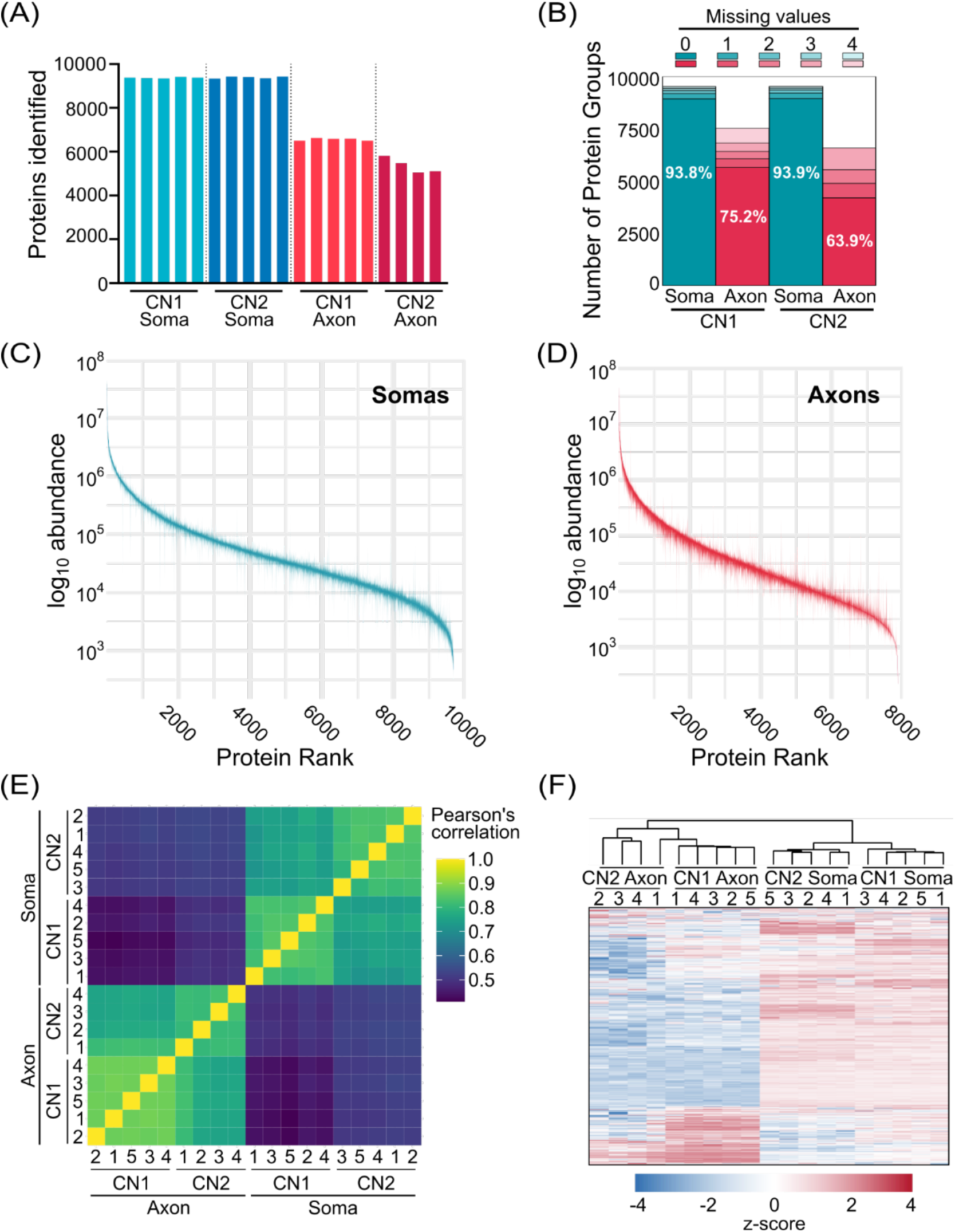
Deep proteomic profiling of human iPSC-derived cortical neurons (CN) harvested from microfluidic devices. **A**, total number of protein groups identified in axonal (Axon) and somatodendritic (Soma) compartments from two CN lines (CN1/CN2). Each bar represents one biological replicate (n = 5; except CN2 Axon, n = 4). **B**, data completeness across replicates for CN1/CN2 Axon and Soma samples showing total number of protein groups identified in each sample type, stratified by number of missing values. Percentages represent proteins present in all replicates with no missing values. **C-D**, protein rank plots for Soma (**C**) and Axon (**D**) samples from highest to lowest median log_10_ abundance with interquartile ranges indicated. **E**, sample correlation matrix showing proteomic similarity by Pearson’s correlation of protein abundances. Samples from the same compartment showed the strongest correlation. **F**, Hierarchical clustering of protein abundances shows compartment-specific enrichment patterns and clustering.

These results demonstrate the power of this ultrafast, ultrasensitive approach for characterising low-input proteomes. As expected, the somatodendritic fraction, which contains axons in addition to somata and dendrites, yielded a broader proteome than the axonal fraction, which contains only axons. Data completeness across biological replicates was very high for somatodendritic samples (94% of proteins detected in all replicates), but lower for axonal samples (75% for CN1; 64% for CN2) (Fig. 2B). This may reflect the challenge of analysing axonal samples yielding only ∼50 ng of peptide input but may also be due to intrinsic variability in axonal growth across cultures. Of note, the CN1 line grew longer axons than the CN2 line, which likely explains the lower number of proteins identified in CN2 axonal samples. Nonetheless, the consistent quantification of more than 5,000 proteins across biological replicates of axonal proteomes demonstrates that deep and reproducible proteome coverage can be achieved from limited axonal material harvested from MFDs.

Rank–abundance profiles demonstrated comparable dynamic range between compartments, with a global shift toward lower intensities in axonal samples consistent with reduced total protein mass (Fig. 2, C-D). Within these profiles, ubiquitously expressed housekeeping proteins such as GAPDH were abundant in both compartments, while nuclear and transcription-related proteins were enriched in the somatodendritic fraction and proteins associated with cytoskeletal regulation and axonal function were preferentially detected in the axonal fraction (Supplemental Table S1). Across biological replicates, protein intensities were highly reproducible within compartments (Fig. 2E). Pairwise correlation analyses showed strong soma–soma (mean Pearson*r = 0.96*) and axon–axon (mean Pearson*r = 0.92*) correlation coefficients, while soma–axon correlations were consistently lower (mean Pearson*r = 0.67*), indicating robust compartment-dependent proteome organisation. Consistent with this, hierarchical clustering segregated samples into distinct somatodendritic and axonal branches (Fig. 2F).

In summary, our Orbitrap Astral DIA-MS approach provides, to our knowledge, the deepest human axonal proteome measured to date. This depth is achieved with excellent reproducibility of quantification across samples, making the method ideally suited for comparative quantitative analysis of the axonal proteome.

### Compartmentalised neuron proteomics identifies a distinct axonal proteome

Principal component analysis (PCA) revealed clear separation of axonal and somatodendritic compartments, with PC1 (57.4%) distinguishing the two compartments and PC2 (17%) capturing variation between independent iPSC lines within each compartment (Fig. 3A). This suggests that subcellular compartment is the primary source of proteomic variation in the dataset. As expected, due to reduced axonal growth, the CN2 axonal samples showed larger variation between replicates. UpSet plot analysis determined that most proteins (6,286) were detected across both compartments in both cell lines (Fig. 3B), consistent with the somatodendritic compartment containing whole neurons, including axons. In line with this, only 13 proteins were detected exclusively in axonal samples (Supplemental Table S2).

**Fig 3.**
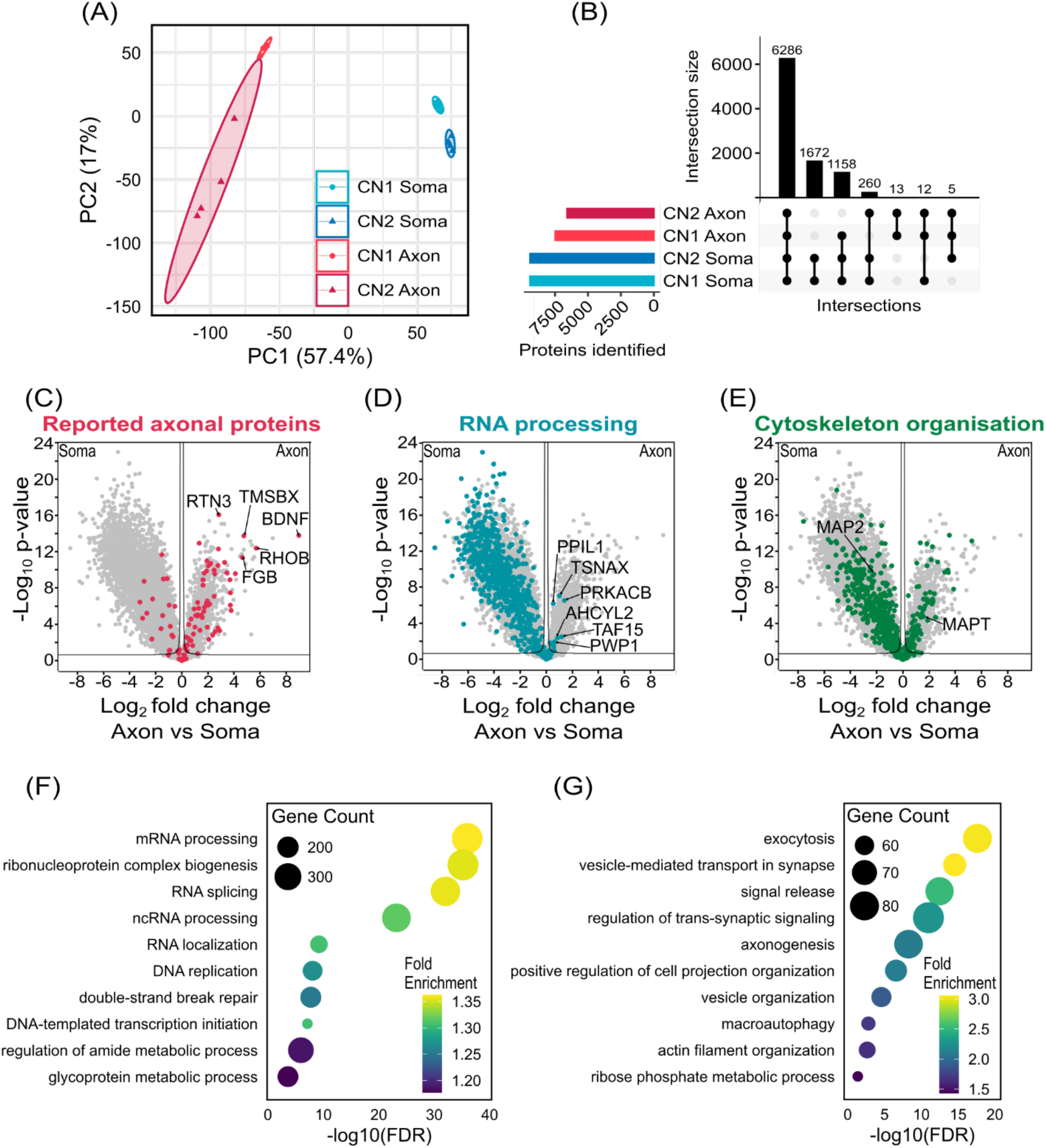
Comparative analysis of axonal versus somatodendritic proteomes. **A**, principal component analysis (PCA) plot for axonal (Axon) and somatodendritic (Soma) samples from two human iPSC-derived cortical neuron lines (CN1/2). **B**, UpSet plot showing overlap of identified proteins between sample groups after filtering for proteins identified in at least 70% of samples for one sample type. Intersections with no overlapping proteins are not plotted. **C–E**, volcano plots showing differential protein abundance between axonal and somatodendritic compartments. 9,406 proteins were quantified across both compartments and data were analysed with a two-tailed Student’s *t*-test (volcano lines indicate the significance threshold with FDR = 0.05, s0 = 0.1). Proteins are coloured according to (**C**) previously reported axonal proteins from Cavarischia-Rega et al.^24^ or gene ontology terms for (**D**) RNA processing (GO:0006396) and (**E**) Cytoskeleton organisation (GO:0007010). Highly enriched axonal proteins are annotated in **C** as well as axonal RNA processing proteins **D**. MAP2 and MAPT are highlighted in **E** to illustrate effective compartmentalisation of dendrites and axons respectively. Further GO terms are annotated in Supplemental Figure S3. Analyses without imputation are also provided in Supplemental Figure S4. **F-G**, gene ontology analysis of significantly enriched proteins in (**F**) somatodendritic and (**G**) axonal compartments, using significance thresholds from **C-E**.

To identify compartment-enriched proteins, we performed differential abundance analysis, identifying 1,250 proteins significantly enriched in the axonal compartment and 6,589 enriched in the somatodendritic compartment (Fig. 3, C-E). Of note, we detected 94 out of 127 proteins previously reported as axon-enriched in human iPSC-derived dopaminergic neurons^24^, with the majority (60) also enriched in the axonal compartment in our analysis (Fig. 3C). This highlights the robustness of the microfluidic approach while also demonstrating the increased depth achieved by our Astral DIA-MS approach. Axon-enriched proteins included neurofilament subunits and synaptic vesicle proteins, confirming robust enrichment of axonal proteins in the distal compartment (Supplemental Fig. S3, A-B). Conversely, somatodendritic-enriched proteins included RNA processing factors (Fig. 3D), as well as nuclear, chromosome, and translation associated proteins, reflecting the concentration of transcriptional machinery in the cell body (Supplemental Fig. S3, C-D). Cytoskeletal components were detected in both compartments consistent with their ubiquitous roles throughout the neuron, with known compartmentalised proteins such as the primarily dendritic MAP2 and the axon restricted Tau (MAPT) segregating accordingly (Fig. 3E).

Gene Ontology (GO) analysis further confirmed functional differences: somatodendritic-enriched proteins mapped to nuclear organisation, transcriptional regulation and RNA metabolism (Fig. 3F), while axon-enriched proteins mapped to vesicle-mediated transport, exocytosis and synaptic vesicle dynamics (Fig. 3G), processes central to axonal transport and presynaptic function.

Together, these results demonstrate that compartmentalised proteomics resolves a functionally distinct axonal proteome and establish a high-confidence resource of axon-enriched proteins (Supplemental Table S3).

### Axonal enrichment analysis of cortical and lower motor neurons reveals a core axonal proteome

Having established our methodology for deep axonal proteomics, we next asked whether the workflow could resolve compartmentalised proteome organisation across distinct neural subtypes. For this, we compared two independently generated iPSC neuronal models representing cortical and lower motor neurons: the cortical neurons (CN1), enriched for deep-layer projection neuron identity, described above, and lower motor neuron-like spinal motor neurons (MN1). CN1 neurons were produced using a neurotrophic factor–guided cortical differentiation protocol that generates predominantly glutamatergic deep-layer projection neurons, alongside a proportion of GABAergic interneurons, as is typical of such protocols^29,30^. MN1 neurons were derived from iPSCs using transcription factor–driven motor neuron programming (*NGN2, ISL1,* LHX3) as previously described^28^. These models therefore differ in both donor genetic background and differentiation strategy, providing an opportunity to identify conserved versus subtype-specific features of the axonal proteome.

Immunocytochemistry confirmed the expected molecular identity of each model, with CN1 cultures expressing layer V/VI corticospinal projection neuron markers (TBR1, CTIP2) and MN1 cultures expressing canonical lower motor neuron markers (ChAT, HB9) (Fig. 4A). Heatmaps of predefined marker sets further validated neuronal subtype identity at the proteome level, with motor neuron identity markers (Fig. 4B) and neurotransmitter subtypes (Fig. 4C) segregating CN1 and MN1 samples according to upper versus lower motor neuron and glutamatergic/GABAergic versus cholinergic profiles, respectively. PCA across all CN1 and MN1 samples (Fig. 4D) revealed that compartment identity (axonal vs somatodendritic) explained the largest fraction of variance (PC1, 52.5%), whereas neuronal subtype (CN1 vs MN1) separated along PC2 (17.4%), indicating conservation of axonal proteome characteristics across differing neuronal subtypes.

**Fig 4.**
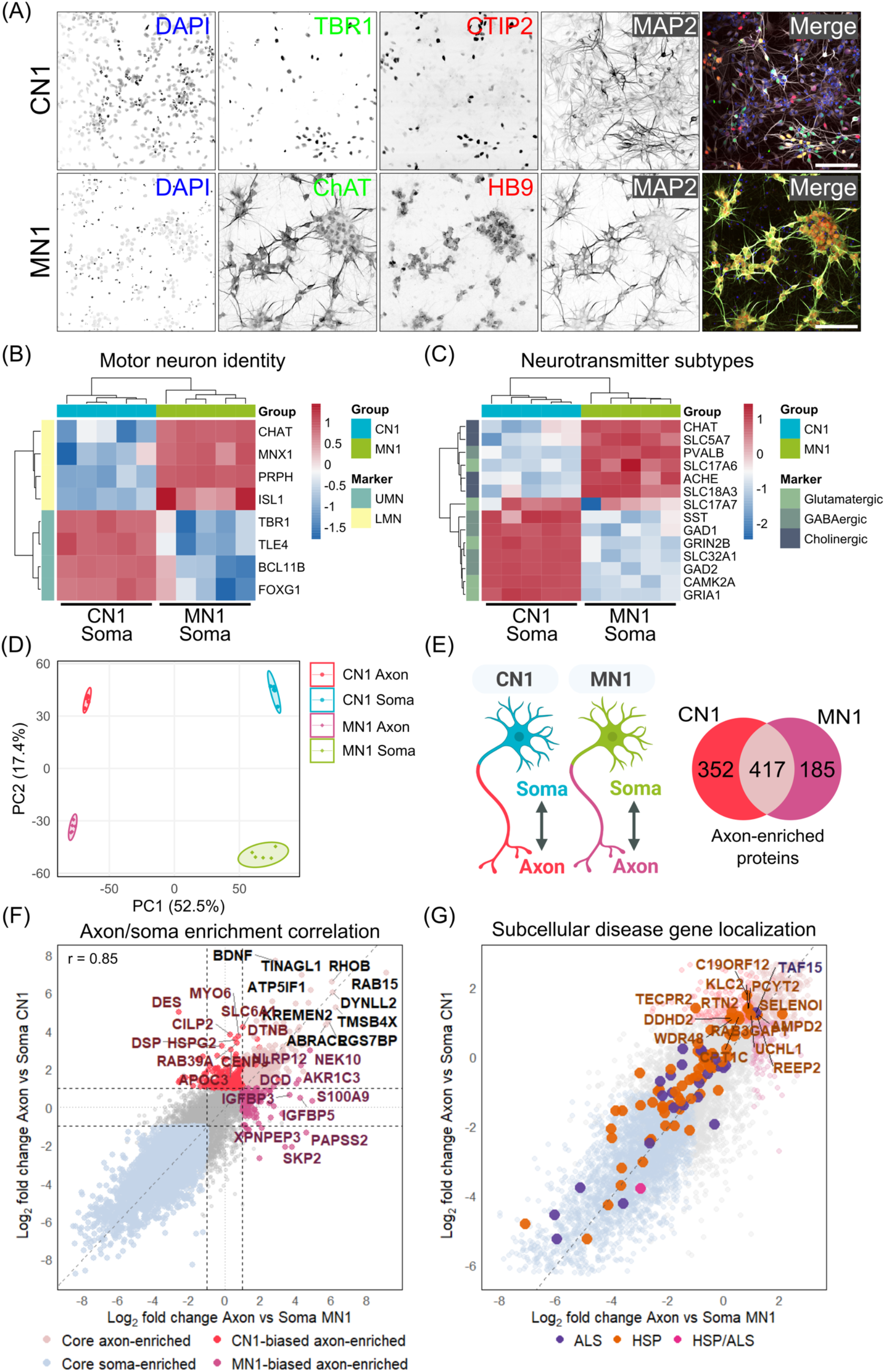
Comparative proteomics analysis of iPSC-derived CN1 and MN1. **A,** Immunocytochemistry of iPSC-derived cortical neurons (CN1; DAPI, TBR1, CTIP2, MAP2) and spinal lower motor neurons (MN1; DAPI, ChAT, HB9, MAP2). Scale bar, 100 µm. **B, C,** Heatmaps of curated marker proteins demonstrating distinct subtype-specific expression patterns, including upper and lower motor neuron identity markers and neurotransmitter-related proteins. **D,** PCA plot of median-normalized proteomes showing primary separation by subcellular compartment (soma vs axon; PC1, 52.5%) and secondary separation by motor neuron subtype (CN1 vs MN1; PC2, 17.4%). **E,** Venn diagram of significantly axon-enriched proteins (log₂FC > 1, p < 0.05, S_0_ = 0.1) in CN1 (769) and MN1 (602), identifying 417 conserved core axonal proteins. **F,** Scatter plot comparing axon-to-soma log₂ fold-changes between MN1 (x-axis) and CN1 (y-axis) for the 8785 proteins quantified in both models. Proteins are coloured by enrichment category: core axon-enriched, core soma-enriched, CN1- and MN1-biased axon-enriched. Top-ranked proteins in axon-enriched categories are labelled. The dashed diagonal indicates perfect correlation; dashed horizontal and vertical lines denote significance thresholds (|log₂FC| = 1). Pearson correlation coefficient (r) is shown. **G,** Enlarged view of panel F highlighting motor neuron disease-associated proteins mapped onto the global proteome distribution, including proteins linked to hereditary spastic paraplegia (HSP, orange), amyotrophic lateral sclerosis (ALS, purple), and a shared HSP/ALS association (pink).

To test this hypothesis, we performed differential abundance analysis to identify axon-enriched proteins for each neuronal model, using more stringent statistical cutoffs than those applied above (FDR = 0.05, s0 = 0.1, log2 fold change > 1). The majority of axon-enriched proteins were shared between models, revealing a core set of 417 proteins constituting a putative conserved axonal proteome across neuronal subtypes (Fig. 4E). Gene Ontology analysis of these core axon-enriched proteins revealed strong enrichment for biological processes related to synaptic vesicle cycling and neurotransmitter release, alongside terms related to axonogenesis and developmental growth regulation (Supplemental Fig. S5). Cellular compartment annotation further supported this profile, highlighting transport vesicles, synaptic membranes, and distal axon structures as the primary localisations of the conserved proteome. Together, these analyses indicate conserved functional pathways in axons across neuronal subtypes, with presynaptic vesicle trafficking machinery being a central feature. Among the most highly axon-enriched proteins in both models were BDNF, a key neurotrophin involved in axonal growth and synaptic plasticity^41^, as well as regulators of vesicle trafficking and cytoskeletal dynamics such as RAB15, DYNLL2, and TMSB4X^42–44^. Enrichment values (log2 fold change axon vs. soma) were highly consistent between CN1 and MN1 (Pearson*r* = 0.85), with only a small subset of proteins enriched in opposite compartments between models (Fig. 4F). Alongside the conserved axonal core, smaller subsets of proteins showed CN1- or MN1-biased axonal enrichment. These proteins were significantly axon-enriched in one model but did not cross the significance threshold in the other.

Finally, to demonstrate the potential of our high sensitivity axonal proteomics workflow for investigating axonal pathology in neurodegenerative diseases, we projected HSP- and ALS-associated proteins onto the axon–soma enrichment space (Fig. 4G). This showed broad coverage of motor neuron disease–implicated proteins in our compartment-resolved dataset, which included 100 of 112 known HSP and ALS-associated genes (89%)^6,45,46^. Disease-associated proteins spanned the full range of enrichment scores, with 15 of 100 classified as axon-enriched, predominantly with a CN1 bias (10/15), 35 as core soma-enriched and the remainder showing no clear compartment preference. This enrichment map provides a quantitative, compartment-resolved reference for assessing how disease-linked mutations can reshape compartmental proteome organisation. For example, to test whether a perturbation shifts proteins between axonal and somatodendritic compartments or selectively disrupts the axon-enriched proteome of a specific neuron type. This framework enables interpretation of disease-associated changes in both protein abundance and subcellular localisation.

### Comparative axonal proteomics reveals differences between cortical and lower motor neurons

Axonal enrichment analysis demonstrated global similarities in protein compartmentalisation between cortical and lower motor neurons. However, the strong axon-soma differences may mask more nuanced differences between the axons of the two models. Therefore, we next performed direct comparative proteomic analysis between neuronal subtypes for each compartment.

Within each compartment, most proteins were identified in both neuron subtypes (5,851 in axons; 9,717 in soma). Despite this, differential abundance analysis identified thousands of differentially expressed proteins (Fig. 5, C-D), suggesting significant differences in the expression profiles of axonal and somatodendritic compartments between the two lines. Of note, of the 3,700 proteins differentially expressed within the axonal compartment, 1,107 were uniquely detected as differentially expressed in axons and not in the somatodendritic compartment, highlighting the importance of compartment-specific analysis (Fig. 5C). As expected, PCA confirmed robust separation of CN1 and MN1 along PC1 in both axonal (66.1% variance) and somatodendritic (62.8% variance) fractions (Fig. 5, E-F). However, the PCA loading plots showed this separation to be driven by a distinct set of proteins in each compartment (Fig. 5, G-H), indicating compartment-specific sources of proteomic variation. In the axon, the highest ranked MN1-associated proteins include SLC18A3 (aka VAChT, vesicular acetylcholine transporter), ChAT (Choline acetyltransferase), SNCG (gamma-synuclein), and SYT2 (Synaptotagmin-2), which are all expected to be enriched in lower motor neuron axons. On the other hand, the highest ranked CN1-associated proteins in axons include several markers of GABAergic cortical interneurons (SLC32A1, SLC6A1, GAD1/2, LAMP5). While this may seem unexpected given that the CN1 differentiation protocol is optimised to generate glutamatergic cortical neurons, co-differentiation of GABAergic interneurons is commonly observed in such protocols^47^. This illustrates how deep axonal proteomics can characterise neuronal models in fine detail.

**Fig 5.**
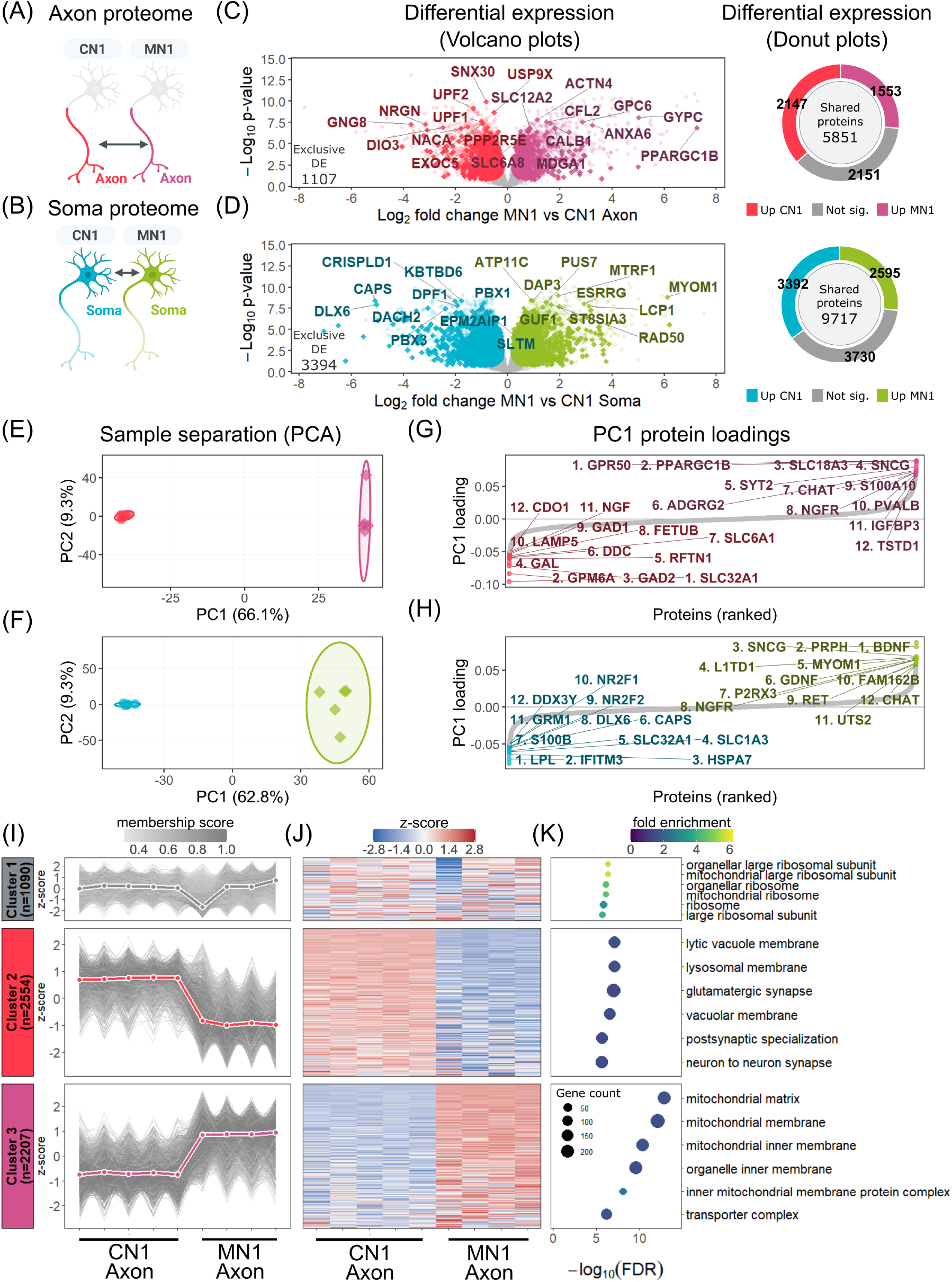
Axonal proteome profiling distinguishes CN1 and MN1 distal axons. Comparative proteomic analysis of CN1 and MN1 axon (top) and soma (bottom) compartments. **A, B,** Schematic of the compartmentalised culture system highlighting the subcellular compartment subjected to proteomic analysis. **C, D,** Volcano plots displaying log₂ fold change versus −log₁₀ p-value for differential protein abundance between MN1 and CN1 in axon (C) and somatodendritic (D) compartments. All significantly enriched proteins are coloured according to their direction of regulation, with CN1-enriched (red in C; blue in D) or MN1-enriched (purple in C; green in D); grey points are non-significant. Numbers in the lower left of each volcano denote counts of compartment-exclusive DEPs, shown as intensely coloured diamonds, whereas shared DEPs are shown as faint circles. The 10 most positively and 10 most negatively enriched proteins are labelled. Donut plots (right of each volcano) summarise the total quantified proteome in each compartment, classified as CN1-enriched, MN1-enriched, or non-significant; centre values indicate the shared detectable proteome. **E, F,** PCA of protein intensities in axon (E) and somatodendritic (F) compartments showing separation of CN1 and MN1 samples along PC1. **G, H**, PC1 loading plots for axon (G) and soma (H) identifying the highest-ranking proteins contributing to sample separation, with the top 12 CN1- and MN1-associated proteins labelled and ranked. **I-K,** Mfuzz fuzzy c-means clustering (c = 3) of the shared axonal proteome (5,851 proteins) partitions proteins into three clusters based on abundance patterns across CN1 and MN1 axon replicates. For each cluster, line plots (I) show z-score–normalized protein abundance trajectories coloured by membership score; heatmaps (J) display per-protein z-scores across all axonal replicates; and dot plots (K) show Gene Ontology Cellular Component (GOCC) enrichment of core cluster members (membership ≥ 0.7; −log₁₀ FDR on the x-axis; dot size scaled to gene count; colour indicates fold enrichment). Cluster 1 (n = 1,090) contains proteins with broadly similar abundance between CN1 and MN1 axons. Cluster 2 (n = 2,554) contains CN1-enriched proteins. Cluster 3 (n = 2,207) contains MN1-enriched proteins. Significance was assessed by permutation-based FDR (q < 0.05) for volcano plots and Benjamini–Hochberg correction for GO enrichment.

Next, we employed Mfuzz soft clustering^37^ to uncover coordinated expression patterns across the shared axonal proteome in an unsupervised manner (Fig. 5I). Unlike hard clustering approaches, this approach assigns each protein a graded membership score across clusters, permitting partial membership and accommodating continuous differences in protein abundance between CN1 and MN1 within a largely shared axonal proteome^38^. Using a three-cluster solution, proteins partitioned into Cluster 1 (similar abundance between CN1 and MN1 axons), Cluster 2 and Cluster 3 (enriched in CN1 and MN1 axons, respectively). Corresponding heatmaps (Fig. 5J) revealed consistent within-cluster patterns across biological replicates, with hierarchical clustering grouping CN1 and MN1 samples separately, indicating robust and reproducible expression signatures. To functionally annotate each cluster, Gene Ontology enrichment analysis was performed on core cluster members (membership ≥ 0.7; Fig. 5K). The CN1-biased Cluster 2 (n = 2,554) was enriched for proteins associated with glutamatergic synapse, as expected for cortical projection neurons, but also for lysosome-related terms such as lytic vacuole and lysosomal membrane (Fig. 5K). In contrast, the MN1-biased Cluster 3 (n = 2,207) was enriched for mitochondrial terms, such as mitochondrial matrix and mitochondrial membrane, while the evenly distributed Cluster 1 was dominated by terms associated with ribosomes. GO Biological Process and KEGG pathway analyses reinforced these functional distinctions, with the CN1-biased Cluster 2 enriched for lysosomal transport, endosomal trafficking and autophagy, and the MN1-biased Cluster 3 enriched for oxoacid metabolism, energy derivation and mitochondrial membrane organisation (Supplemental Fig. S7). Notably, several of these axon-enriched terms, including lysosomal membrane, proteasome-associated complexes and endomembrane trafficking components were not detected when the same enrichment analysis was applied to the somatodendritic proteome (Supplemental Fig. S8). This demonstrates that compartment-resolved analysis reveals biological processes that are obscured in whole-cell measurements.

In conclusion, these results demonstrate that deep, quantitative axonal proteomics reveals both conserved and subtype-specific molecular features of distal axons that are obscured in conventional whole-cell analyses, underscoring its value for dissecting axonal biology in disease-relevant models.

## Discussion

### A deep and scalable workflow for human axonal proteomics

Understanding the molecular composition of the axon is fundamental to explaining how these specialised cellular compartments sustain function over vast distances and why they appear selectively vulnerable in neurodegeneration. Here, we present a deep, compartment-resolved proteomic map of human axonal and somatodendritic compartments from iPSC-derived cortical neurons, achieved by coupling microfluidic axon isolation with DIA-MS on an Orbitrap Astral platform. We quantified an average of 6,029 proteins in the axonal compartment and 9,382 in the somatodendritic compartment (Fig. 2A), representing the deepest human axonal proteome reported to date. This provides both a workflow and a rich resource for dissecting compartment-specific proteostasis in neuronal health and disease.

The depth achieved by our study substantially advances the state of the art for microfluidic axonal proteomics. In human iPSC-derived dopaminergic neurons, Cavarischia-Rega et al. identified ∼500 protein groups in the axonal compartment using a SILAC (Stable Isotope Labelling with Amino acids in Cell culture) DDA-MS approach, of which 127 proteins were either exclusively detected in the axon or axon-enriched^24^. The authors’ elegant use of dynamic SILAC allowed them to identify 154 proteins that were locally synthesised in the axon before transport to the soma. However, the combination of DDA-MS, which is susceptible to undersampling due to stochastic precursor selection, and SILAC, which splits peptide signal across channels, constrains proteomic depth achievable from the ∼50 ng protein that is recoverable from microfluidic axonal compartments. Highlighting the increased depth achievable with label-free DIA-MS, Zare et al. leveraged DIA-PASEF on a timsTOF Pro 2 to identify approximately 2,400 axonal proteins from primary mouse motor neurons^25^. Notably, our study used a 21-minute LC gradient, whereas Zare et al. used a 120-minute gradient. Thus, our workflow achieves a ∼2.5-fold increase in identifications in approximately one sixth of the time. This gain in depth despite a substantially shorter gradient reflects the combined sensitivity and acquisition speed of narrow-window DIA on the Orbitrap Astral, which together maximise proteome coverage from the limited ion flux available from low-input axonal material, without requiring extended separation to achieve sufficient depth^48,49^. Importantly, we identified 94 of the 127 proteins previously identified as axon-enriched in human iPSC-derived dopaminergic neurons (Cavarischia-Rega et al., 2024), of which the majority (60) were also axon-enriched in our dataset (Fig. 3C). This provides orthogonal validation of their axonal localisation, while our data also demonstrate the substantially increased depth afforded by our Astral DIA-MS approach.

### A distinct and functionally specialised axonal proteome

A defining feature of our dataset is the robust and reproducible proteomic distinction between axonal and somatodendritic compartments. Although very few proteins were identified exclusively in axons, as the somatodendritic-enriched compartment also contains axons, differential abundance analysis identified 1,250 axon-enriched proteins, representing an approximately tenfold increase relative to the previous human dataset^24^. Gene ontology analysis showed that axon-enriched proteins map to pathways including vesicle-mediated transport, exocytosis, synaptic vesicle dynamics, axonogenesis, and synaptic signalling, while somatodendritic-enriched proteins were associated with DNA replication and repair, transcriptional regulation, and RNA metabolism (Fig. 3, F-G) – consistent with the known biology of each compartment. The enrichment of synaptic vesicle machinery (e.g., SYP, SYN1, SYN2, SNAP25, VAMP2, SV2A) confirms presynaptic specialisation of the distal axon, while the axonal enrichment of neurofilament subunits (NFL, NFM) reflects their central roles in maintaining axonal architecture, cytoskeletal spacing and organelle positioning (Supplemental Fig. S3, A-B)^50^. These functional signatures align with*in vivo* proximity labelling data from mouse dopaminergic axons^51^, extending these observations across species and neuronal subtypes and substantially expanding the catalogue of axon-enriched proteins.

### Conserved and subtype-specific features of the axonal proteome

By profiling two independently derived iPSC neuronal models – upper motor neuron-like cortical projection neurons (CN1) and lower motor neuron-like spinal motor neurons (MN1) – we were able to assess conserved and subtype-specific features of the human axonal proteome. PCA revealed that cellular compartment (axonal vs. somatodendritic) was the dominant source of proteomic variance (PC1, 52.5%), exceeding the contribution of neuronal subtype (PC2, 17.4%) (Fig. 4D). Axonal enrichment values between the two cell lines were highly correlated (Pearson r = 0.85) (Fig. 4F), demonstrating the robustness of our approach and supporting conserved compartmentalisation of the proteome across neuron subtypes. Stringent differential abundance analysis identified a core set of 417 proteins significantly axon-enriched in both models (Fig. 4E), including neurofilaments, molecular motors, synaptic vesicle proteins, and mitochondrial components.

These proteins represent fundamental effectors of axonal maintenance regardless of subtype identity, donor genetic background, or differentiation strategy. Alongside this conserved core, unsupervised soft clustering of the shared axonal proteome revealed subtype-specific functional signatures (Fig. 5, I-K). Lower motor neuron-biased axonal proteins were enriched for mitochondrial matrix and membrane components, alongside KEGG pathways including pyruvate metabolism, branched-chain amino acid degradation, and porphyrin metabolism (Fig. 5K; Supplemental Fig. S7) – pathways that collectively support mitochondrial TCA cycle activity and electron transport chain maintenance. This is consistent with the well-established high energetic demands of cholinergic motor neurons^52,53^, though functional validation will be required to determine whether these proteomic differences reflect genuine differences in axonal mitochondrial activity between the two neuronal subtypes. By contrast, cortical neuron-biased axonal proteins were unexpectedly enriched for lysosomal categories including lytic vacuole and lysosomal membrane, lysosomal transport and autophagy (Fig. 5K; Supplemental Fig. S7). Notably, these categories were not enriched in the corresponding somatodendritic analysis (Supplemental Fig. S8), indicating that lysosomal enrichment is an axon-specific feature of cortical projection neurons that is diluted below detection in whole-cell lysates. This provides a clear illustration of the value of compartment-resolved proteomics for uncovering biologically meaningful signals masked by bulk measurement. Though the functional basis of this enrichment is not yet clear, active degradative lysosomes are continuously delivered to distal axons to maintain local degradative capacity^54,55^. Whether cortical projection neuron axons depend more heavily on lysosomal machinery than lower motor neuron axons remains an open question that warrants future investigation.

Intriguingly, these subtype-specific axonal signatures are consistent with known patterns of selective vulnerability in motor neuron disease. Lysosomal and autophagic dysfunction is a prominent pathogenic theme in HSP, with mutations in several HSP-associated genes directly disrupting endolysosomal trafficking, autophagosome biogenesis and autophagic lysosome reformation^56,57^. The relative enrichment of lysosomal machinery in cortical neuron axons observed here may therefore reflect a heightened dependence on local degradative clearance, providing a potential framework for understanding why disruption of these pathways preferentially causes upper motor neuron degeneration. Conversely, the mitochondrial enrichment in lower motor neuron axons is consistent with their well-documented vulnerability to energetic stress in ALS^52^. Whether these proteomic differences contribute to the selective vulnerability of each neuronal subtype in disease remains to be determined. Furthermore, as our two neuronal models differ in both donor genetic background and differentiation strategy, observed subtype-specific proteomic differences cannot be unambiguously attributed to neuronal identity alone, making replication in isogenic lines important for mechanistic follow-up.

### Implications for axon-selective vulnerability in motor neuron disease

To further illustrate the disease relevance of our axonal proteomics pipeline, we mapped HSP- and ALS-associated proteins onto the compartment-resolved enrichment space (Fig. 4G). Both HSP and ALS are characterised by axon pathology, in which distal axons and neuromuscular junctions are prominently affected^6,9,11^, indicating that disease-relevant molecular events may be concentrated within the axonal compartment. Strikingly, however, most HSP- and ALS-associated proteins did not show axonal enrichment in healthy neurons (Supplemental Fig. S6), despite the predominantly axonal pathology of these diseases. This indicates that the selective vulnerability of axons to perturbation of these genes is unlikely to reflect preferential localisation in the axonal compartment but may instead arise from downstream consequences that affect axonal function, for example, disruption of axonal transport, local proteostasis, or energy metabolism. Nevertheless, a subset of disease-associated proteins was axon-enriched, including KLC2, which mediates anterograde axonal transport as part of the kinesin-1 complex and has been shown to localise within axons in a punctate pattern^58^. The ER-shaping proteins RTN2 and REEP2, essential drivers of tubular ER formation^59,60^, were also enriched in the axonal compartment, consistent with the predominantly tubular structure of axonal ER – in contrast to somatic ER, which comprises both sheets and tubules. PCYT2 and SELENOI, enzymes of the Kennedy pathway for phosphatidylethanolamine biosynthesis that are mutated in HSP^61,62^, were also axon-enriched, representing a novel observation that may have implications for understanding how disruption of membrane lipid homeostasis leads to axonal pathology.

Our quantitative compartment-resolved reference proteome now enables the application of our workflow to iPSC-based disease models, where genetic perturbations can be interrogated not only for their effects on overall protein abundance, but also for their spatial consequences within the neuron. This distinction, which is inaccessible to whole-cell proteomics, is likely to be important for understanding compartment-specific pathology in motor neuron diseases.

## Acknowledgement

This study has been supported by the German Ministry of Education and Research (BMBF grant 01GM1905A, 01GM2209F to LS, SH), the Tom Wahlig foundation (to AD, SH) and the Academy of Medical Sciences Springboard scheme [SBF009\1160], funded by the Academy of Medical Sciences, the Wellcome Trust, the Department for Science, Innovation and Technology, the British Heart Foundation and Diabetes UK (to AD). JS, RS, and BK were supported by core funds from the Technische Universität Dresden. Schematics were created using BioRender (Sauter, C. (2026) https://BioRender.com/522127i), licensed under CC BY 4.0. We thank Matthias Mann (Max Planck Institute of Biochemistry) for generously providing access to the Orbitrap Astral mass spectrometer and for his support of this project.

## Author contributions

Conceptualization: CMS, AD, SH

Data curation: CMS, ZS, AD

Formal analysis: CMS, ZS

Funding acquisition: LS, AD, SH

Investigation: CMS, MKo, VA, MKr, RS, BK

Methodology: CMS, ZS, VA, AD, SH

Resources: JS, LS

Supervision: JS, LS, AD, SH

Visualization: CMS, ZS

Writing – original draft: CMS, ZS, AD

Writing – review and editing: SH, AD

## Declaration of generative AI and AI-assisted technologies in the manuscript preparation process

During the preparation of this work, the authors used ChatGPT (OpenAI) and Claude (Anthropic) to assist with the development and troubleshooting of R scripts used for data visualisation. After using these tools, the authors reviewed, tested and edited all code as needed and take full responsibility for the content of the published article.

