## Supplementary Data for "Deep axonal proteomics of human iPSC-derived neurons by microfluidic separation and DIA-MS"

### Supplemental figures

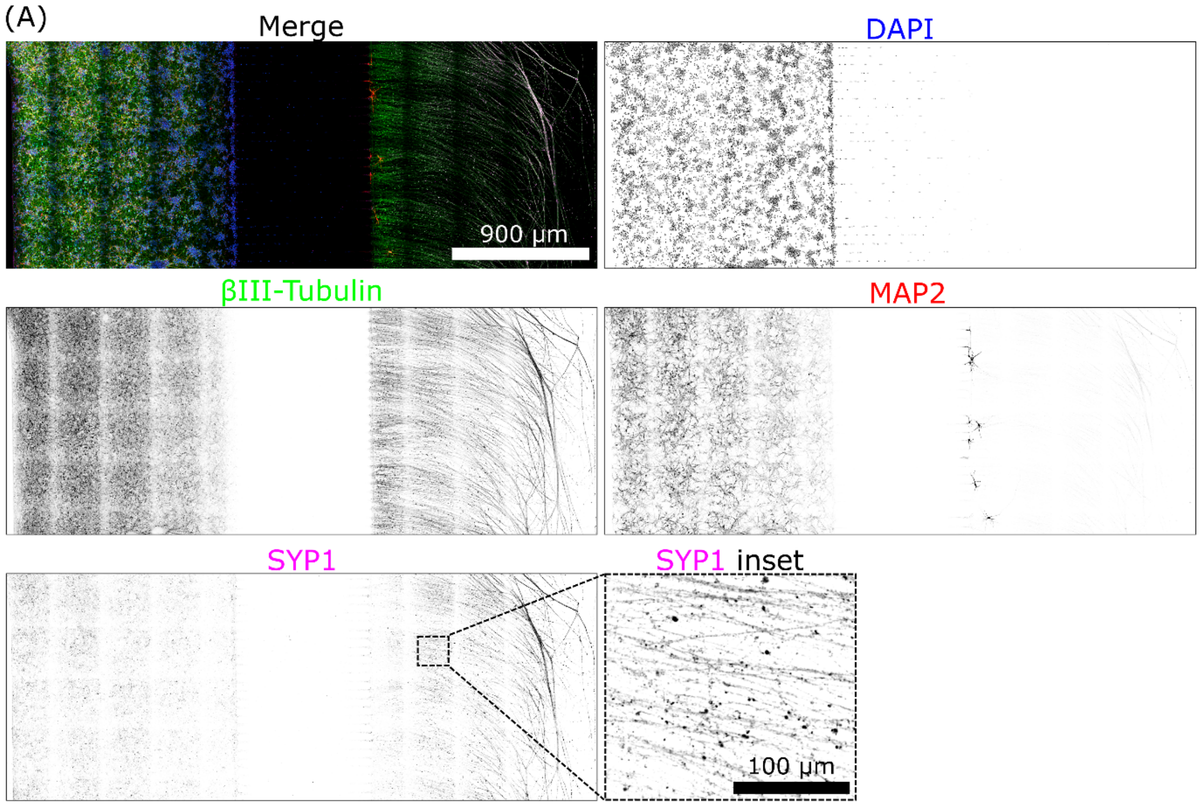


**Supplemental Figure S1. Immunocytochemical validation of neuronal compartment identity. A,** Representative immunocytochemistry showing DAPI, βIII-tubulin, MAP2, and Synaptophysin-1 (SYP1) staining and the corresponding merge. βIII-tubulin labels neuronal processes, MAP2 marks somatodendritic compartments, and SYP1 identifies presynaptic vesicle clusters. SYP1-positive puncta are observed along βIII-tubulin–positive projections in regions morphologically consistent with axons. The boxed region in the merge is shown at higher magnification in the SYP1 inset, highlighting punctate staining characteristic of presynaptic structures.Scale bars: 900 µm (merge), 100 µm (SYP1 inset).

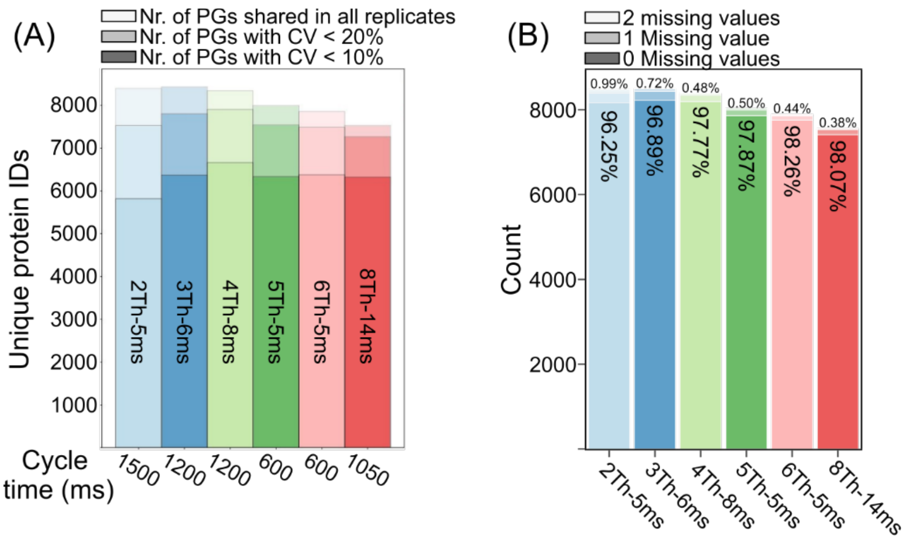


**Supplemental Figure S2. Optimization of DIA isolation window width and injection time.** Six methods with varying isolation window widths (2–8 Th) and injection times (5–14 ms) were compared using 50 ng soma samples. **A,** Bars show protein groups shared across triplicates (light), with CV < 20% (medium), and CV < 10% (dark). **B,** Bars show protein groups across triplicate stratified by missing values 0 (dark), 1 (medium) or 2 (light) missing values. Percentages within the bars represents the proportion of proteins identified in all replicates (0 missing values) and above the bar represent proteins only in one repeat (1 missing value).

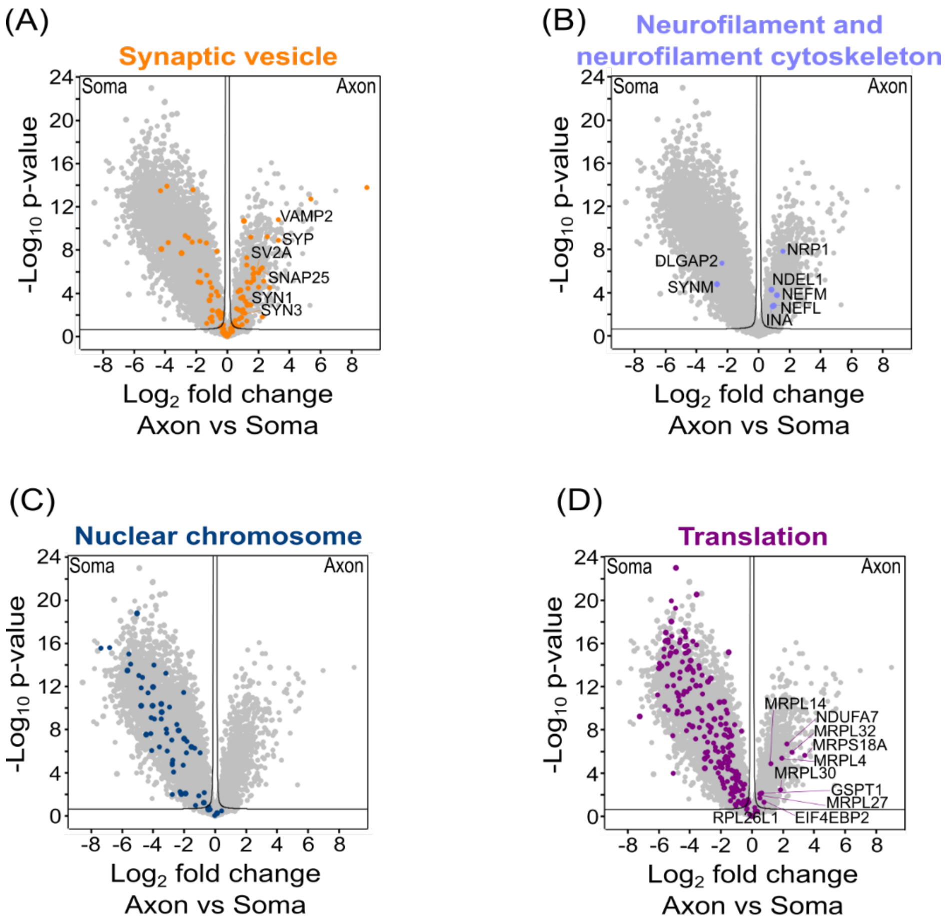


**Supplemental Figure S3. GO-term analysis of CN1 and CN2 axons vs somas.** Volcano plots of compartment-enriched proteins from CN1 and CN2 cell lines. Axes show $\log_2$ fold-change (axon/soma) and -log(p-value) from Student's t-test (s0=0.1, FDR=0.05). Synaptic vesicle (GO:0008021; **A**), neurofilament and neurofilament cytoskeleton (GO:0005883 and GO:0060053; **B**) nuclear chromosome (GO:0000228; **C**) and translation (GO:0006412; **D**) are presented with outlying proteins and proteins of note annotated.

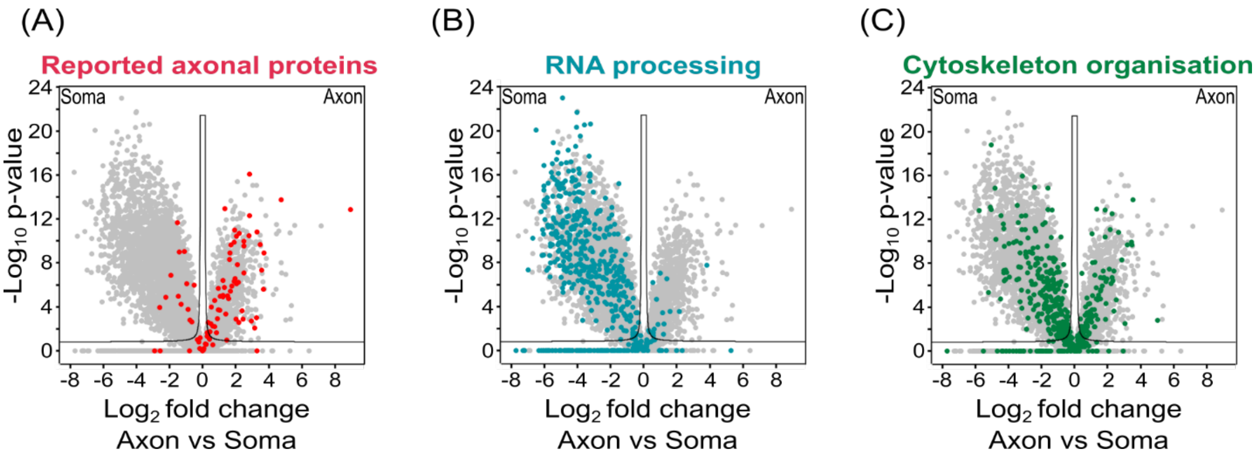


**Supplemental Figure S4. Differential expression analysis of axonal versus somatodendritic proteomes without imputation.** Volcano plots showing differential protein abundance between axonal and somatodendritic compartments as described in Figure 3. Proteins were filtered to those identified in at least 70% of samples for one sample type and data were analysed with a two-tailed Student's t-test (volcano lines indicate the significance threshold with FDR = 0.05, s0 = 0.1). No data imputation was performed. Proteins are coloured according to (**A**) previously reported axonal proteins from Cavarischia-Rega et al.[24] or gene ontology terms for (**B**) RNA processing (GO:0006396) and (**C**) Cytoskeleton organisation (GO:0007010). Results are consistent with those shown in Figure 3.

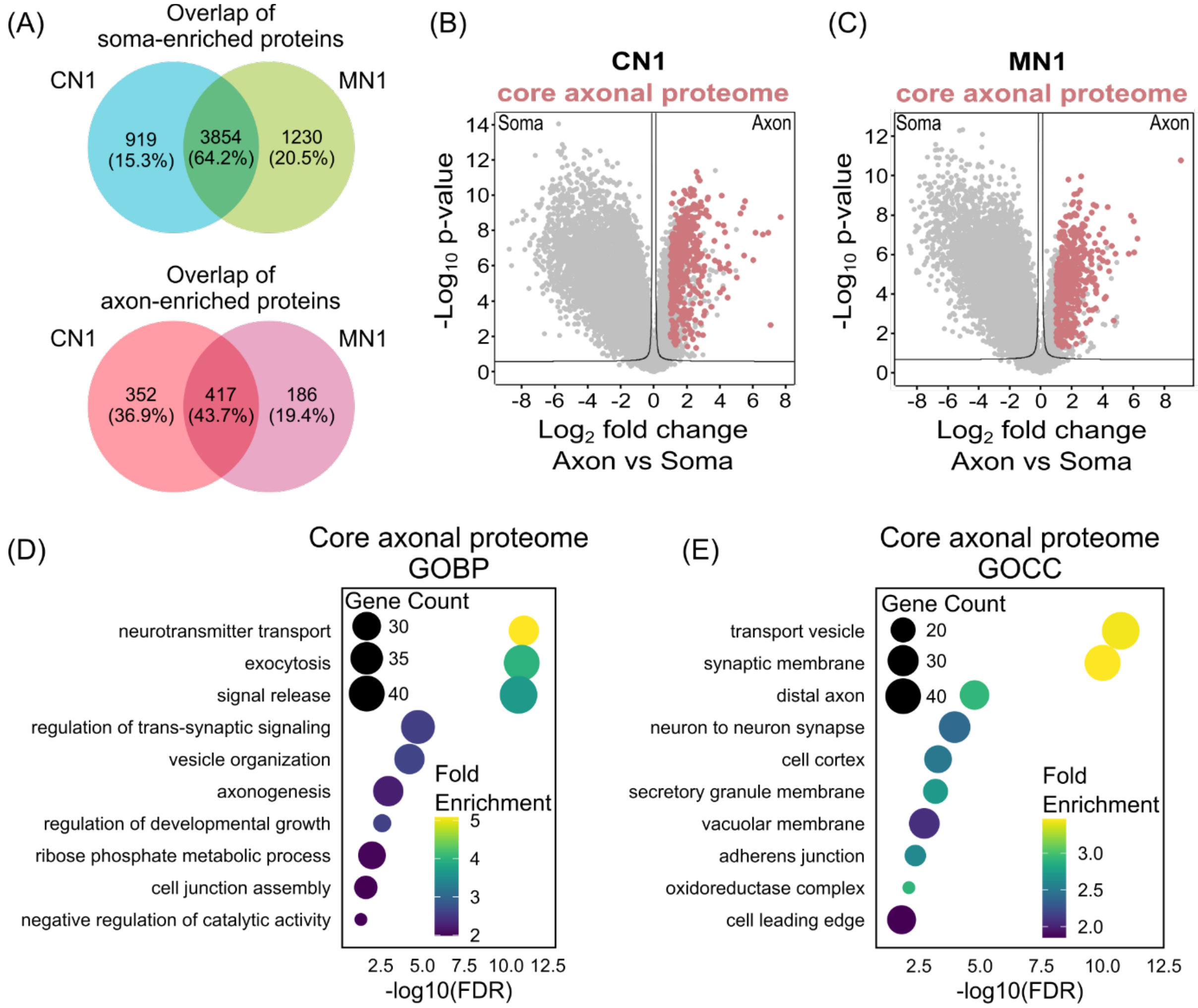


**Supplemental Figure S5. Core axonal proteome enrichment in CN1 and MN1 cell lines. A,** Venn diagram showing the overlap between somatodendritic- and axon-enriched proteins ($log_2FC > 1$, $p < 0.05$, $S0 = 0.1$) identified in CN1 and MN1 datasets. **B, C,** Volcano plots showing differential protein abundance between axonal and somatodendritic compartments for CN1 **(B)** and MN1 **(C)** cell lines. Proteins significantly enriched in axonal compartments for both CN1 and MN1 are highlighted in red. **D, E,** Gene ontology analysis of the shared core axonal proteome, showing enriched biological processes **(D)** and cellular compartments **(E)**.

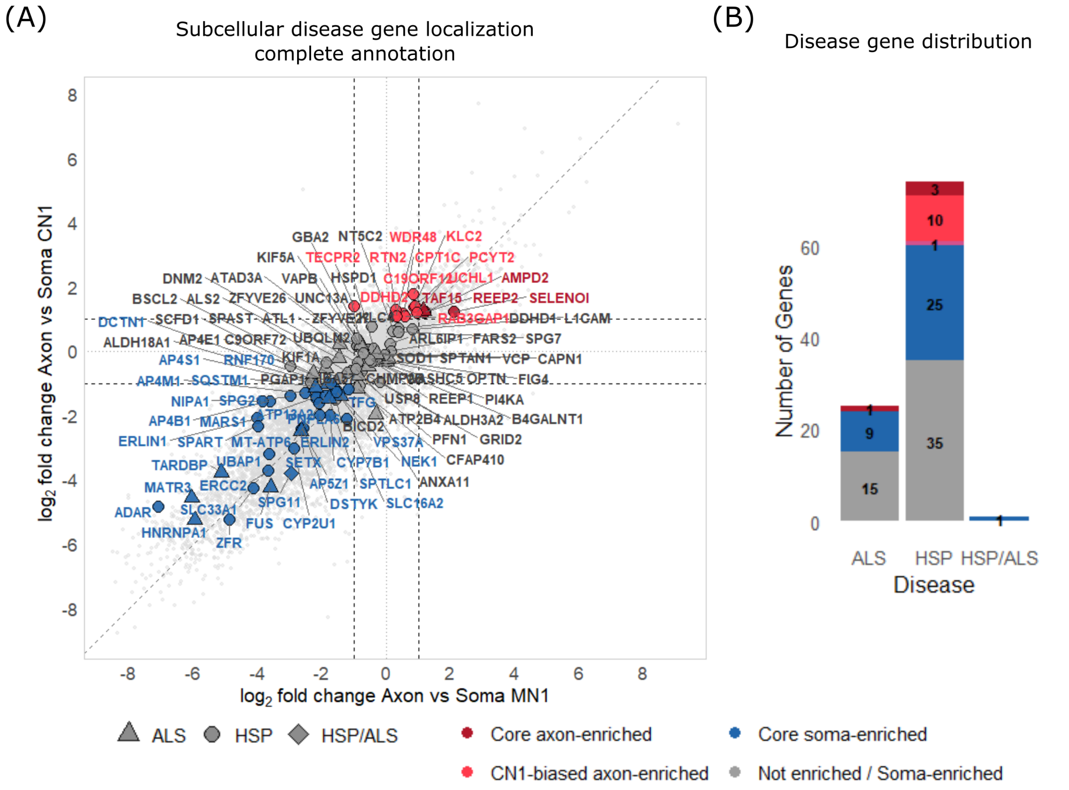


**Supplemental Figure S6. Subcellular localization of HSP/ALS disease genes. A,** Correlation of axon–soma $\log_2$ fold change between CN1 and MN1 datasets. Disease-associated genes (HSP, ALS, HSP/ALS) are highlighted and classified into the four categories: core axon-enriched, neuron-type–biased axon-enriched, core soma-enriched, or not enriched / Soma-enriched ($|\log_2 FC| > 1$; $-\log_{10} p > 1.3$). The dashed diagonal indicates concordant subcellular localization across neuronal types. D, Proportion of disease-associated genes (HSP, ALS, HSP/ALS) falling into each compartment category.

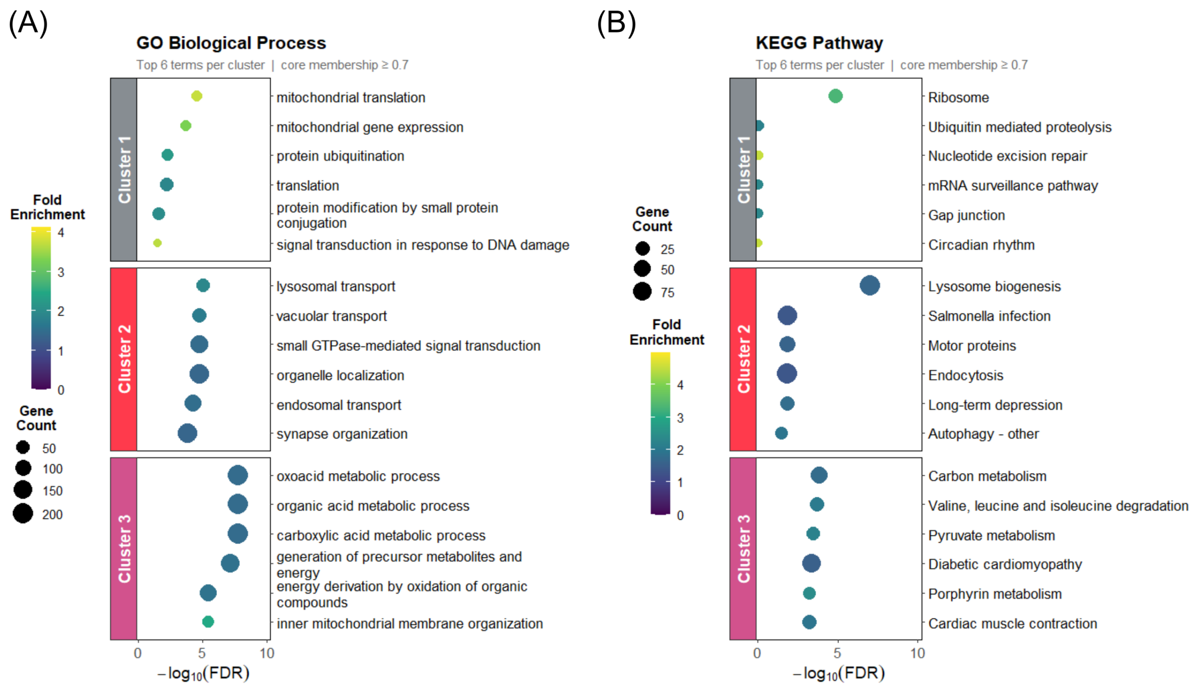


**Supplemental Figure S7. GO Biological Process and KEGG pathway enrichment of Mfuzz clusters from the shared axonal proteome. A, B,** GOBP (A) and KEGG pathway (B) enrichment analysis of core cluster members (membership > 0.7) from fuzzy c-means clustering of the shared axonal proteome (see Fig. 5I-K). Dot plots show the top 6 enriched terms per cluster; the x-axis represents $-\log_{10}$(FDR), dot size is scale to gene count and colour indicates fold enrichment. Significance was assessed using the Benjamini-Hochberg method (adjusted $p < 0.05$).

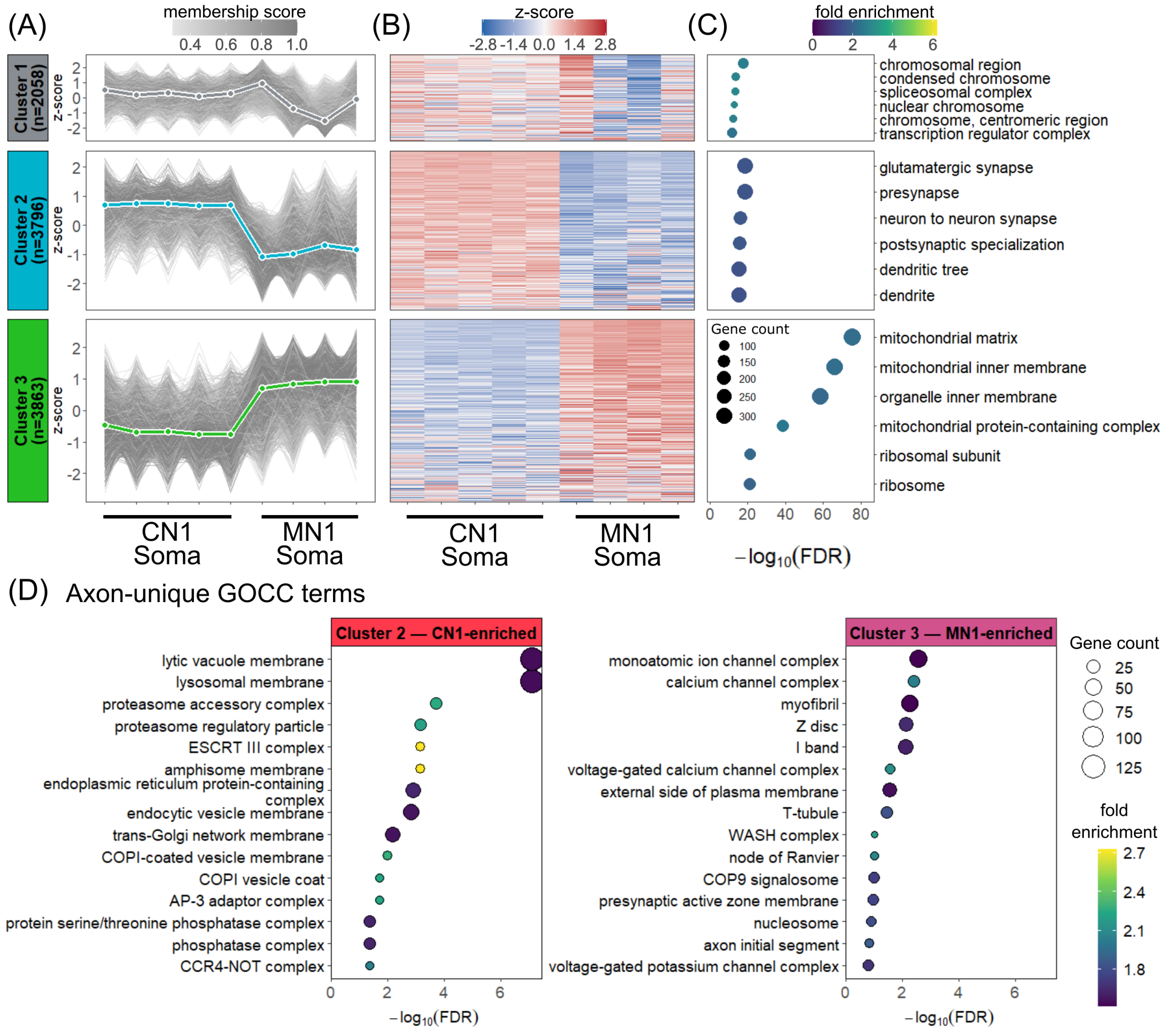


**Supplemental Figure S8. Mfuzz clustering of the soma proteome and axon-unique GOCC enrichment. A-C,** Mfuzz fuzzy c-means clustering (c = 3) of the shared soma proteome (9,717), presented as in Fig. 5 I-K. Line plots (A), heatmaps (B) and GOCC enrichment dot plots (C) are shown for Cluster 1 (2,058), Cluster 2 (3,796) and Cluster 3 (3,863). **D,** GOCC terms uniquely enriched in the axonal but not the somatodendritic proteome. Enrichment analysis was performed independently on each compartment (p.adjust < 0.05, fold enrichment > 1.5), yielding 82 and 166 significant terms for the axon and soma CN1-enriched cluster, and 59 and 70 for the MN1-enriched cluster, respectively. Cross-referencing identified 30 axon-unique, 114 soma-unique and 52 shared terms for CN1-enriched proteins, and 28 axon-unique, 39 soma-unique and 31 shared for MN1-enriched proteins. Dot plots show the top 15 axon-exclusive terms for CN1-enriched (Cluster 2, left) and MN1-enriched (Cluster 3, right) proteins; dot size corresponds to gene count and fill colour indicates fold enrichment.

**Supplemental Table S1. Most abundant proteins in CN1 and CN2 axon and soma samples.** The top 30 most abundant protein groups in CN1 and CN2 axon and soma samples are list below. Protein groups present in each top 30 are highlighted in bold. Abundance values represent normalised median $log_2$ intensities for each compartment after filtering for common contaminants.

| **Rank** | **Most abundant soma protein groups** | **Median log2 intensity in somas** | **Most abundant axon protein groups** | **Median log2 intensity in axons** |
|---|---|---|---|---|
| 1 | **GAPDH** | 25.21392 | **GAPDH** | 25.44103 |
| 2 | H4C1 | 25.0355 | **BASP1** | 25.3922 |
| 3 | INS;INS-IGF2 | 24.97311 | **DPYSL2** | 24.61817 |
| 4 | H1-4 | 24.95253 | **TUBB2B** | 24.26781 |
| 5 | H2AC18;H2AC20 | 24.78288 | MAPT | 23.80841 |
| 6 | **CFL1** | 24.38419 | **CRMP1** | 23.69418 |
| 7 | **TUBB2B** | 23.9659 | GDI1 | 23.19254 |
| 8 | H1-5 | 23.87011 | **DPYSL3** | 23.16026 |
| 9 | **HSP90AA1** | 23.75739 | PRDX5 | 23.09691 |
| 10 | **CRMP1** | 23.71986 | **CKB** | 23.06821 |
| 11 | **DPYSL2** | 23.61852 | MIF | 23.06534 |
| 12 | HSP90AB1 | 23.53931 | GNB1 | 22.97973 |
| 13 | TRAP1 | 23.46712 | MDH1 | 22.97619 |
| 14 | **DPYSL3** | 23.37315 | **GNAO1** | 22.95274 |

| | | | | |
|---|---|---|---|---|
| 15 | HNRNPA2B1 | 23.28601 | **HSP90AA1** | 22.93199 |
| 16 | HNRNPA1 | 23.21198 | RAB15 | 22.86735 |
| 17 | H2BC18 | 23.11901 | TPI1 | 22.82625 |
| 18 | **HSPA8** | 23.06775 | UCHL1 | 22.82499 |
| 19 | STMN1 | 22.93621 | CALM1;CALM2;CALM3 | 22.81514 |
| 20 | MARCKS | 22.86899 | **CFL1** | 22.79112 |
| 21 | **UBB;UBC** | 22.74575 | **HSPA6** | 22.706 |
| 22 | **GNAO1** | 22.65887 | ATP1A3 | 22.70592 |
| 23 | **HSPA6** | 22.55772 | **YWHAG** | 22.68132 |
| 24 | **BASP1** | 22.54115 | INA | 22.6511 |
| 25 | **YWHAG** | 22.52817 | **UBB;UBC** | 22.57556 |
| 26 | PKM | 22.46715 | HSP90AB1 | 22.57443 |
| 27 | **CKB** | 22.40875 | DCX | 22.54856 |
| 28 | PEBP1 | 22.38625 | **HSPA8** | 22.46839 |
| 29 | HSPE1 | 22.37745 | FSCN1 | 22.43977 |
| 30 | **ATP5F1B** | 22.36254 | **ATP5F1B** | 22.41056 |
